# Predicting species invasiveness with genomic data: is Genomic Offset related to establishment probability?

**DOI:** 10.1101/2024.02.20.581132

**Authors:** Louise Camus, Mathieu Gautier, Simon Boitard

## Abstract

Predicting the risk of establishment and spread of populations outside their native range represents a major challenge in evolutionary biology. Various methods have recently been developed to estimate population (mal)adaptation to a new environment with genomic data via so-called Genomic Offset (GO) statistics. These approaches are particularly promising for studying invasive species, but have still rarely been used in this context. Here, we evaluated the relationship between GO and the establishment probability of a population in a new environment using both in silico and empirical data. First, we designed invasion simulations to evaluate the ability to predict establishment probability of two GO computation methods (Geometric GO and Gradient Forest) under several conditions. Additionally, we aimed to evaluate the interpretability of absolute Geometric GO values, which theoretically represent the adaptive genetic distance between populations from distinct environments. Second, utilizing public real data from the crop pest species *Bactrocera tryoni*, a fruit fly native from Northern Australia, we computed GO between “source” populations and a diverse range of locations within invaded areas. This practical application of GO within the context of a biological invasion underscores its potential in providing insights and guiding recommendations for future invasion risk assessment. Overall, our results suggest that GO statistics represent good predictors of the establishment probability and may thus inform invasion risk, although the influence of several factors on prediction performance (e.g. propagule pressure or admixture) will need further investigation.

## 1 Introduction

Predicting how natural species adapt to environmental change is crucial in evolutionary biology. This knowledge not only furthers our understanding of evolutionary processes involved in adaptation but also holds practical implications, particularly in the current context of global changes. At the within-species level, the variation of abiotic and biotic conditions over a species distribution range favor the evolution of locally adapted populations, which has been documented in a wide diversity of taxonomic groups (Wadgymar *et al*., 2022). This dynamic process, rooted in genetic variations, intricately shapes the fitness-related traits of organisms in diverse habitats. Thus, characterizing the genetic underpinnings of local adaptation over varied habitats as snapshots in space can inform on how populations might respond or adapt over time to local environmental alterations. With the advent and democratization of high-throughput sequencing technologies, such a “space-for-time substitution” approach has thus been recently introduced in the population genomics field to forecast the potential impact of changing conditions on species’ vulnerability (Capblancq *et al*., 2020b; Rellstab *et al*., 2021).

In practice, these population-based approaches rely on Genome-Environment Association (GEA) methods to detect associations between the frequencies of adaptive genetic variants and environmental or phenotypic covariables. The GEA step also enables modeling the structure (or composition) of genetic diversity across populations and discerning which adaptive genetic variants and extrinsic variables may play a role in adaptation (Ruegg *et al*., 2018; Bogaerts-Márquez *et al*., 2021; Capblancq *et al*., 2020a; Ingvarsson and Bernhardsson, 2020). Subsequently, these statistical associations can be used to predict the optimal theoretical genetic composition (i.e., frequencies for different adaptive genetic variants) providing the highest fitness in a new given environment. The difference between the optimal genetic makeup in a new environment (according to the GEA modeling) and that of a population of interest has been referred to as Genomic Offset (GO) (Rellstab *et al*., 2021; Gain *et al*., 2023). It is aimed at quantifying maladaptation risk: higher GO indicates greater risk of mismatch between population genetic composition and the new environment. Alternatively, GO measures can be viewed as weighted environmental distances based on covariables characterizing the environment of the populations, weighted according to their relative impact on the adaptive genetic composition of the populations (Gain *et al*., 2023).

In recent years, GO approaches have been widely adopted in the field of conservation biology, being employed across a diverse array of taxa to estimate populations vulnerability to climate change to inform future management action (Ruegg *et al*., 2018; Rhoné *et al*., 2020; Morgan *et al*., 2020; Zhang *et al*., 2023a; Capblancq *et al*., 2020a; Borrell *et al*., 2020). These methods indeed allow accounting for population local adaptation within species, which is not achievable through conventional Species Distribution Modelling (SDM) methods that assume niche uniformity. Another application field where GO measures could prove useful is for the study of invasive populations. Indeed, global trade and climate change amplify the need to develop strategies allowing to predict future biological invasions (Gallien *et al*., 2010) and to mitigate their negative impacts (Hulme, 2017, 2021), especially for species of agricultural interest, e.g., crop pests which represent a significant threat to global food security (Bruce, 2010; Bradshaw *et al*., 2016). In this context, GO measures could help predict the optimal regions for population establishment, taking into account intraspecific local adaptation.

Over the past few years, several methods have been proposed to compute GO. Among them, two widely used are the Risk Of Non-Adaptedness or RONA (Rellstab *et al*., 2016) and Redundancy Analysis or RDA (Capblancq and Forester, 2021), which model a linear relationship between allele frequencies and extrinsic covariates. Another recent linear method, known as Geometric GO (gGO) has shown particularly promising performance (Gain *et al*., 2023). Geometric GO relies on estimates of regression coefficients between population allele frequencies and environmental variables using Latent Factor Mixed Modelling or LFMM (Frichot *et al*., 2013), i.e. the effect sizes of each environmental covariable on the variation of allele frequencies, while accounting for neutral population structure through the simultaneous estimation of latent factors (i.e., confounding factors). Under certain conditions, gGO is strictly equivalent to GO computed with RDA (Gain *et al*., 2023). Conversely, two alternative methods have been early proposed to estimate GO in the pioneering study by Fitzpatrick and Keller (2015) relying on Gradient Forest (GF) algorithm (Ellis *et al*., 2012)) and Generalized Dissimilarity Modelling (GDM) approach (Ferrier *et al*., 2007) that accommodate non-linear relationships between allele frequencies and covariates. They rely on turnover curves that describe the rate of genetic change along a gradient of environmental values, but do not account directly for the confounding effects of neutral population structure. The GF method, based on the machine learning random forest approach (Breiman, 2001), has been particularly used in many recent studies (Ruegg *et al*., 2018; Adam *et al*., 2022; Lachmuth *et al*., 2023; Zhang *et al*., 2023b). Beyond the modeling differences summarized above, all GO approaches assume that i) the populations adapt to their environment through pre-existing variants (rather than *de novo* mutations); ii) the genome-environment relationship remains constant over time for future predictions (“space-for-time” hypothesis); and iii) the populations studied are already adapted to their current environment (Capblancq *et al*., 2020b; Rellstab *et al*., 2021). Several studies recently evaluated GO-based methods by comparing predicted maladaptation (quantified with GO) against fitness-related traits. This was done through in silico simulations (Ĺáruson *et al*., 2022; Gain *et al*., 2023; Lotterhos, 2023) or by studying populations in common gardens (Rhoné *et al*., 2020; Archambeau, 2022; Fitzpatrick *et al*., 2021). Encouragingly, these studies often found that higher predicted maladaptation aligned with reduced realized fitness. However, none of these studies considered the specific situation of biological invasions, although GO measures are beginning to be applied within this framework (Chen *et al*., 2023, 2021).

In response to this gap, we here propose an evaluation of GO measures in the context of biological invasions. A primary objective of our study is to assess the predictive performance of GO, and we therefore evaluate the correlation between several GO measures and invasive population establishment probability. We further propose some methodological innovations including i) a novel approach to compute gGO (gGO BayPass) based on the BayPass GEA model (Gautier, 2015); ii) an improvement in the computational efficiency of the Gradient Forest package to accommodate larger datasets; and iii) an evaluation of interpretation of the absolute value of gGO, most GO metrics being relative and not directly tied to measurable aspects thereby complicating their biological interpretation across different datasets. Finally, for illustrative purpose and to exemplify the practical application of GO in the context of biological invasion, we also analyzed the data recently published by Popa-Báez *et al*. (2020) regarding *Bactrocera tryoni*, a tropical invasive fruit pest fly, native from Australia.

## 2 Materials and Methods

### 2.1 Simulation study

To evaluate the relevance of GO in the context of biological invasion, we simulated the evolutionary dynamics of populations under biological invasion scenarios using SLiM v4.0.1 (Messer, 2013). In a first phase, we simulated 25 populations, each consisting of 1,000 individuals, within a native area under a Wright-Fisher model including spatial structure and non-uniform selection constraints in order to produce genetic patterns of local adaptation. In this native area, each population was associated with two environmental optima (with values ranging from –1 to 1), to which individuals adapt through QTNs (Quantitative Trait Nucleotide). In a second phase, individuals from a given source population in the native area were randomly selected to invade a new environment, also characterized by given values for the two environmental optima.

The simulation framework in the native area was inspired by the work of Ĺaruson *et al*. (2022), who evaluated the correlation between several GO measures and population fitness. However, because one of the primary objectives of our study was to examine GO in the context of biological invasion, we adapted the simulations, especially in the invaded area, to more closely match life history traits of invasive species, such as an important number of offspring, short generation time and overlapping generations (Sakai *et al*., 2001; Zhao *et al*., 2023) (Supplementary Note 1). While these characteristics are shared by many invasive species, the specific parameter values considered here were informed by literature focusing on invasive insect biology (Supplementary Note 1), which represents a major concern in agriculture, as arthropods largely contribute to emerging alien species (Seebens *et al*., 2021; Bradshaw *et al*., 2016).

#### 2.1.1 Genome and trait architecture

The simulated genome totaled 250 cM and consisted of five chromosomes of 50 cM each modeled as a segment of 5×10^5^ sites with a per-site per generation recombination rate of 10*^−^*^5^. Mutations affecting the phenotypes (QTNs) were simulated on the first four chromosomes at a rate of 2.5 x 10*^−^*^8^ with effect sizes (in units of standard phenotypic deviations) independently sampled from a Gaussian distribution with a mean of 0 and a standard deviation of 0.1. Each QTN was characterized by two effect sizes contributing to two distinct phenotypes. For each phenotype, the effect sizes of all the QTNs present in an individual genome were summed to form its phenotypic value. The two phenotypes determined the individual fitness, which reached its maximum value of 1 when the two phenotypes matched the two environmental optima and decreased when the distance between the phenotypes and the environmental optima increased (Supplementary Note 1). As fitness represents an individual probability of being selected as a parent for the next generation in the native area, this mechanism favored the spread of locally advantageous mutations. Note that given the distribution of QTNs effect sizes, a given phenotypic value between –1 and 1 could be obtained through multiple combinations of approximately fifteen mutations per individual, implying a high degree of genetic redundancy.

In addition to the QTNs simulated directly during SLiM forward approach, neutral mutations were added afterward using the python packages msprime (v1.2.0) and pyslim (v1.0.3) according to the *recapitation and overlay* procedure described by Haller *et al*. (2019), at a rate of 1.0 x 10*^−^*^7^ per generation per site. Mutations with a MAF *<* 1% were filtered out using the program VCFtools (v 0.1.16) (Danecek *et al*., 2011).

#### 2.1.2 Simulations design: native area

Evolution in the native area was simulated under a 2D stepping stone model with selection, where slight modifications were made compared to Ĺaruson *et al*. (2022) work (Supplementary Note 1). The native area consisted of 25 populations of 1,000 individuals each, organized in a 5×5 grid that evolved under a Wright-Fisher demographic model with non-overlapping generations and constant population size. Populations exchanged individuals with either high (0.05) or low (0.005) migration rates, resulting in varying degrees of local adaptation. As mentioned above, the grid position was associated with two different values of environmental optima.

To investigate the impact of the adaptive landscape and its relation with neutral genetic structure, three different distributions of the environmental optima were considered (Figure 1). Simulations of the native area included 3,000 SLiM generations, which were divided into the 3 following phases i) an initialization of 1,000 generations without any environmental variation in order to generate the necessary adaptive genetic variation; ii) 1,000 generations with a gradual transition of the environment towards the specified optima value; and iii) 1,000 final generations under the adaptive landscape.

**Figure 1:**
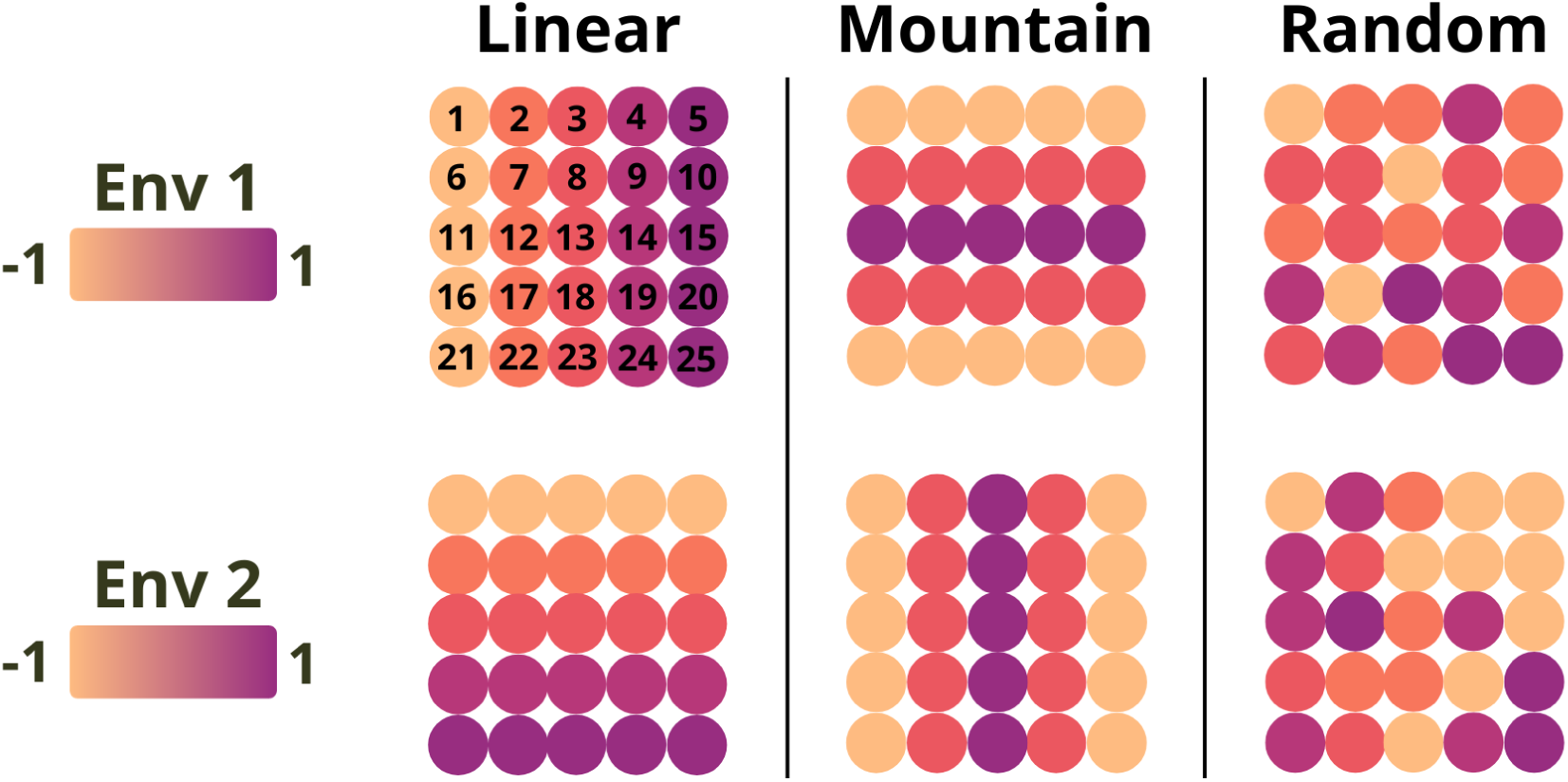
S**c**hematic **representation of the three simulated native environment types**. For each environment type (columns), the two grids represent the optimal values of environmental variables *e*_1_ (top) and *e*_2_ (bottom) for each of the 25 populations. Population indices are indicated on the top left panel and specify the source populations used for invasion (see Simulations design: invaded area).

This procedure and its duration of 3,000 generations were established as sufficient to ensure robust alignment between populations and their environment (Supplementary Note 1 and Figure S1). Ten random replicates of the evolutionary history were simulated for each of the 6 native environ-ment scenarios (3 environment types *×* 2 migration rates). “Mountain” (M) and “Random” (R) environments aimed to reduce environment-demography correlation seen in the “Linear” (L) envi-ronment. The realized genome-wide *F_ST_* across all populations for each native area was computed with the *computeFST* function (*poolfstat* R package, version 2.1.1) with default settings.

#### 2.1.3 Simulations design: invaded area

After 3,000 simulated generations in the native area, a few founder individuals from a given source population were randomly chosen to invade a new environment. In order to test the effect of the founding bottleneck intensity, we considered either 10 or 100 founder individuals. For each native environment scenario, three distinct source populations were selected for invasion based on their native environmental optima *e*_1_ and *e*_2_: i) *e*_1_ = *e*_2_ = *−*1 (“-1/-1”); ii) *e*_1_ = *e*_2_ = 0 (“0/0”); or iii) *e*_1_ = *e*_2_ = 1 (“1/1”). For example, in the case of the L scenario, these three source populations corresponded respectively to the populations 1, 13 and 25 (Figure 1). The source population could then invade nine possible environments, which correspond to the nine combinations of the three possible environmental values (–1, 0 or 1) for *e*_1_ and *e*_2_ in the invaded area.

Considering the distribution pattern of environmental values in the three simulated native area types (L, R and M), it is important to note that multiple populations could correspond to any of the three potential source environments (e.g., population 1, 5, 21 and 25 all corresponding to the –1/-1 source population in the M environment, Figure 1). In this case, a single source population was arbitrarily selected to simulate invasion.

In the invaded area, we relied on the so-called non-Wright Fisher simulation mode of SLiM (Messer, 2013) to allow for population extinction and estimate establishment probabilities. Reproduction and death events were disconnected in the invasive population, allowing in particular for overlapping generations. The fitness of an individual, derived from its QTN genotypes, quantified its probability of surviving to the next simulated time step (up to a maximum age of 3 time steps). Note that for simplicity, we then further assumed that all living individuals in a given time step had the same probability of reproducing.

Each invasion was repeated 250 times, and the establishment probability (EP) in the new envi-ronment was determined by tallying successful establishment events among these repetitions. A population was deemed established if it exceeded 50,000 individuals or persisted for at least 100 time steps. These thresholds were selected based on preliminary tests, which showed that popula-tions meeting these criteria never faced extinction (Supplementary Note 1 and Fig. S2)

### 2.2 GO estimation

Two different types of approaches have been considered to estimate GO, differing in their underlying modeling of the relationship between genomic composition of the populations and their local environment. The first, named Gradient Forest and proposed by Fitzpatrick and Keller (2015) relies on a Random Forest machine learning algorithm, and the resulting GO estimate will be hereafter referred to as GO_gf_. The second was proposed by Gain *et al*. (2023) and is based on a linear regression model to compute the geometric GO (gGO) defined as:

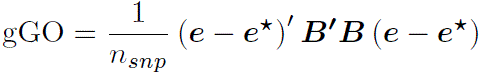

where *n_snp_* is the number of genotyped SNPs; ***e*** and ***e^*^*** are the vectors of *m* environmental covariables values for the two compared environments; and ***B*** is the *n_snp_ × m* matrix of the regression coefficients *β_jk_* of environmental variable *k* on population allele frequencies at SNP *j* as estimated under a GEA linear model. Three different methods were considered to compute gGO differing on how the *β_jk_*’s were estimated. First, we considered the original approach implemented in the *lfmm2* function of the R package *LEA* (v.3.12.2) (Gain and Fraņcois, 2021), where the *β_jk_* were estimated under a Latent Factor Mixed Model (LFMM) (Caye *et al*., 2019). We hereafter refer to the resulting gGO estimator as gGO_lfmm_. For all simulations, we modelled K=2 latent factors since (as expected from the simulation design) these were found sufficient to capture the neutral population structure based on a Principal Component Analysis (PCA) of the genotypes matrix. Alternatively, we considered two estimations of gGO based on *β_jk_*’s estimated under the BayPass Bayesian hier-archical GEA model (Gautier, 2015) which are both implemented in the newly developed R function compute_genetic_offset available from the latest version (2.41) of the software package (Gautier, 2024). One important difference of BayPass compared to LFMM resides on the full modeling (and estimation) of the neutral covariance structure among the population allele frequencies through the so-called **Ω** matrix. First, the standard covariate model was run with default options to estimate **Ω** and the posterior mean of the regression coefficients *β_jk_*’s based on an Importance Sampling (IS) algorithm. The resulting gGO estimator derived from the IS estimates of the *β_jk_*’s is referred to as gGO_is_. We also ran BayPass with option –covmcmc (setting **Ω** to the posterior mean estimated in the first run) to obtain estimates of the *β_jk_*’s as the posterior means from values directly sampled via a Markov Chain Monte Carlo (MCMC) algorithm (instead of relying on the IS approximation). This gGO estimator will be hereafter referred to as gGO_mc_. When considering a single environmental covariate (univariate case), estimates of the regression coefficients were found to be far more accurate with the MCMC than with the IS algorithm (Gautier, 2015). However, it is important to stress that when multiple covariates are analyzed (multivariate case), the MCMC algorithm considers all these covariates jointly (similar to LFMM), while the IS algorithm treats each covariate independently.

Note that other popular methods based on linear modeling to compute GO, namely RONA (Rellstab *et al*., 2016) or RDA-based (Capblancq and Forester, 2021), were not considered here since their properties and relationship with gGO_lfmm_ have been explored in a recent study, and they were shown to be either strictly equivalent (for RDA-based under certain hypotheses) or less accurate (Gain *et al*., 2023). In this previous study, GO_gf_ was also found to generally provide lower prediction accuracy than gGO_lfmm_, at least under the simulation scenario the authors explored. Nevertheless, we kept it in our study for its ability to account for non-linear relationships between allele frequencies and covariates. To estimate GO_gf_, we used a customized version of the original *Gradient Forest* package (v.0.1.32) (Ellis *et al*., 2012) to allow efficient analyses of large number of SNPs (Supplementary Note 2). Note that this optimized version is so far restricted to continuous covariables. To account for neutral population structure, we used as response variable in the GF modeling the residuals of the SNPs allele frequencies obtained after fitting a linear model with K=2 latent factors (Caye *et al*., 2019) as described below (see details in Supplementary Note 2). Finally, following Gain *et al*. (2023), the resulting GO_gf_ estimates were squared to ensure similar scaling than other distance measures.

We also computed the Euclidean distance between the environmental covariables as 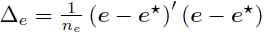 to establish a baseline prediction performance of GO measures that would only include the environmental information but not the genetic one. Indeed, as highlighted by the above gGO definition expression (but less clear with GO_gf_), the GO may directly be interpreted as a weighted environmental distance, whose weights are related to the influence of environmental covariables on the structuring of genetic diversity (quantified by the *B* matrix of regression coefficients in gGO). In other words, a covariable with no impact on genetic diversity (i.e., no SNPs with frequencies associated with its variation) would be highly penalized, and the GO is expected to be null if no SNP is found associated to any covariable. Like GO_gf_, all Euclidean distances were squared.

### 2.3 Evaluation of GO measures

#### 2.3.1 GO to predict establishment probability

To evaluate the predictive accuracy of the different GO measures for establishment probabilities, allele frequencies (for QTNs and neutral markers) in the native area were estimated from 50 randomly sampled individuals per population, keeping only SNPs with a MAF*>*1% to train the GEA models (via GF or linear modeling with LFMM or BayPass).

GO calculations included either all (neutral and QTN) the SNPs or the top 10% overly differentiated SNPs based on the XtX*^*^* statistics estimated with BayPass (Olazcuaga *et al*., 2020; Gautier, 2015). Further, we either considered i) the two causal environmental covariables alone (assuming an unrealistic situation where these would be known); ii) 8 covariables including these 2 causal plus 6 confounding variables; or iii) the first 4 PCs obtained after performing a PCA of these 8 covariables with the R package *ade4* (v1.7-22) (Dray and Dufour, 2007). The 6 confounding covariables consisted of 2 “fake” covariables without any link with the causal variables; two covariables correlated with causal environment 1 (r=0.4 and r=0.8); and 2 variables correlated with causal environment 2 (r=0.4 and r=0.8). The fake variables were randomly generated for each possible environment (25 in the native grid, and 9 potential invaded environments) from a Gaussian distribution so that their correlations with the two causal covariables were close to 0 (ranging between –0.1 and 0.1).

The different GO estimators were then compared based on their Spearman’s correlation R^2^ with the logarithm of the establishment probabilities (log(*p_e_*)) obtained with simulations as detailed above; each R^2^ value was based on a total of 90 observations, arising from the combination of 9 possible invaded environments (for a given source population) and 10 replicates of the native environment scenario (Figure S3). In cases where GO was computed with confounding variables, each GO computation was performed for three distinct random draws of confounding environments and the mean R^2^ value across these three confounding environments was reported. The associations between GO and population fitness, as well as between GO and population growth rate, were investigated using the same approach.

#### 2.3.2 Interpretation of the absolute value of gGO in terms of ***f*_2_**

As demonstrated by Gain *et al*. (2023), gGO values calculated for new environments vary proportionally with fitness logarithms. However, the proportionality coefficient is challenging to compute making it difficult to accurately predict fitness values in a new environment based on estimated gGO. Alternatively, gGO can be interpreted as the expected squared allelic frequency difference across all adaptive loci between two populations, aligning with the definition of the *f*_2_ statistic (Patterson *et al*., 2012). This provides a way to evaluate the accuracy of the absolute value of different GO estimators based on simulated data. We thus compared estimated GO with *f*_2_ for pairs of populations taken from different environments within the native area. The choice of native area populations ensured that allele frequencies were at equilibrium, as assumed in the theoretical prediction of Gain *et al*. (2023).

The gGO calculations were performed between the three potential source populations for invasion and the nine possible combinations of environmental values used to build invaded environments. We here only compared the gGO_lfmm_ and gGO_mc_ estimators since both rely on the same modeling approach consisting of treating all the covariables jointly. The *f*_2_ statistics were estimated using the R package *poolfstat* (version 2.1.1) (Gautier *et al*., 2022), based on the allelic frequencies computed with the genotypes for all the 1,000 individuals of a population. In all cases, we computed Mean Percentage Absolute Error (MAPE) to compare the estimated gGO to their corresponding estimated *f*_2_ (expected to represent the truth).

Two different settings were investigated. We first evaluated the accuracy of gGO under ideal conditions, where the estimators were computed using all causal QTNs; the allele frequencies were computed based on the genotypes for all the 1,000 simulated individuals of a population; and only the two causal environmental variables were used to compute gGO_lfmm_ and gGO_mc_. Second, we evaluated gGO estimates under more realistic conditions, where causal loci are unknown. gGO measures were then computed using QTNs and neutral SNPs and population allele frequencies were obtained from 50 randomly sampled individuals (discarding all SNPs with a MAF*<* 1%). These more ‘realistic’ GO estimates were compared to estimated *f*_2_ computed solely with QTNs or with both QTNs and neutral markers (filtered on MAF), derived from allele frequencies of all individuals within populations. Theoretically, gGO should reasonably predict the *f*_2_ computed solely with QTNs, although a perfect match is not expected due to the exclusion of some QTNs with low MAF. On the other hand, the *f*_2_ computed with both QTNs and neutral markers (MAF-filtered) should be seen as a higher bound that would be reached by a gGO estimation procedure failing to distinguish adaptive and neutral markers.

### 2.4 Bactrocera tryoni case study

#### 2.4.1 Studied populations

We used publicly available data on 28 populations of *Bactrocera tryoni* including 15 native and 13 non-native populations (Figure S13), which were previously analyzed by Parvizi *et al*. (2023) and Popa-Báez *et al*. (2020). The dataset consisted of 6,707 SNPs, which were obtained through Diversity Arrays Technology (DArT) sequencing data for 301 individuals (from 4 to 31 individuals per population).

#### 2.4.2 Environmental data

Environmental data were downloaded from the Chelsa (v2.1, accessed the 27th March 2023) database (Karger *et al*., 2017) using the *dismo* v 1.3.5 (Hijmans *et al*., 2023a) and *raster* v 3.5.15 (Hijmans *et al*., 2023b) R packages. A total of 21 environmental covariables were extracted for each of the studied populations consisting of the averaged values over the period 1981-2010 (at a 30 arc sec resolution) for the 19 commonly used bioclimatic variables, the mean monthly climate moisture index and the mean monthly-near surface wind speed. Indeed, previous works have shown the importance of humidity on fitness-related traits or geographic distribution in *B. tryoni* (Hulthen and Clarke, 2006; Weldon and Taylor, 2010; Sutherst and Yonow, 1998; Dominiak *et al*., 2006). While moisture’s and humidity impact has been established, wind-related variables emerge as potentially influential factors for *B. tryoni*. Wind patterns might affect the presence of dew (Dominiak *et al*., 2006) and could also impact predation dynamics (Dominiak, 2012). To address variable interdependence, we carried out a PCA on all the covariates and retained the first 5 PCs (explaining 95% of the total variance) for GO calculation. To also adopt a similar method as that used by Parvizi *et al*. (2023) for managing variable correlation, we performed the identical analysis but with the six bioclimatic variables they selected (bio 3, bio 5, bio 8, bio 9 and bio 12), showing Pearson correlation coefficients inferior to 0.6.

#### 2.4.3 GO computation

Following the simulation study, we estimated GO_gf_, gGO_lfmm_, gGO_is_ and gGO_mc_ (see above). Bay-Pass analysis was performed here from the allele count file already formatted by Parvizi *et al*. (2023). To ensure comparability across the different estimators in the context of real data including missing genotypes, gGO_lfmm_ was estimated using the allele frequencies obtained from the allele count file, missing data (for 3 different SNPs in 3 different populations) being replaced by the mean of allele frequencies across the 28 populations. For the LFMM analysis (and gGO_lfmm_ estimation), we included K=3 latent factors following Parvizi *et al*. (2023). For the GF analysis, neutral population structure was accounted for by using the residuals of the LFMM analysis as input variable, similar to what was done in the simulation study (Supplementary Note 2).

The different GO estimators were then computed between a “source” population and 12,838,400 positions encompassing an extensive area in Oceania, covering areas that have been invaded or are potentially at risk of invasion. The choice of the source population was based on the estimation of heterozygosities with *poolfstat* package (v.2.2.0). Among the native range, population 1 exhibited the highest heterozygosity (Figure S4). This aligns with *B. tryoni*’s historical records (Popa-Báez *et al*., 2020; Parvizi *et al*., 2023), so this population was selected as the source population to calculate GO.

## 3 Results

### 3.1 GO to predict establishment probability

The primary objective of our simulation study was to assess the predictive capacity of several GO measures in estimating the establishment probability of invasive populations originating from diverse locations within a native area. This evaluation included locally adapted populations across three distinct types of native areas and considered variations in demographic parameters such as migration and the number of invading individuals (see Material and Methods and Figure 1).

Between 1,507 and 1,568 QTNs were simulated under conditions of low migration, while between 1,756 and 1,955 QTNs were obtained for high migration scenarios, as detailed in Table 1. The majority of these QTNs had a low polymorphism level, with only between 44 and 64 (resp. 80 and 109) retained with MAF*>* 1% for the low (resp. high) migration scenario. Mean fitness at the end of the 3,000 simulated generations was lower for populations experiencing higher migration rates. Furthermore, two levels of local adaptation were generated, with low migration exhibiting stronger population differentiation, with *F_ST_* values ranging from 3.4% to 7.2%, in contrast to higher migration scenarios, where *F_ST_* values ranged from 0.34% to 0.46%.

**Table 1:** Information about native area simulations. Means are computed over 10 replicates for each native area, and at the end of the 3,000 simulated generations (see Table S1 for standard deviations that are all two to three orders of magnitude lower).

| Migration rate | Environment type | Mean nb. of QTNs | Mean nb. of QTNs (MAF > 1%) | Mean nb. of neutral mut. (MAF > 1%) | Mean fitness | Mean $F_{ST}$ (in %) |
| --- | --- | --- | --- | --- | --- | --- |
| 0.005 | Linear | 1,507 | 44 | 11,860 | 0.952 | 3.4 |
|  | Mountain | 1,568 | 64 | 11,781 | 0.933 | 4.9 |
|  | Random | 1,535 | 62 | 11,873 | 0.945 | 7.2 |
| 0.05 | Linear | 1,756 | 82 | 11,548 | 0.883 | 0.34 |
|  | Mountain | 1,955 | 109 | 11,504 | 0.807 | 0.40 |
|  | Random | 1,852 | 80 | 11,509 | 0.815 | 0.46 |

We first focus on the results obtained for the low migration scenario (strongest population structuring) with 10 founding individuals in the invaded area. Following Gain *et al*. (2023), we compare Pearson’s correlation of different GO estimates with log(*p_e_*) in order to evaluate their ability to correctly rank the establishment probabilities of a population in different environments. Only the results based on all SNPs (without SNP pre-selection) are presented in the main text; they are depicted in Figure 2.

**Figure 2:**
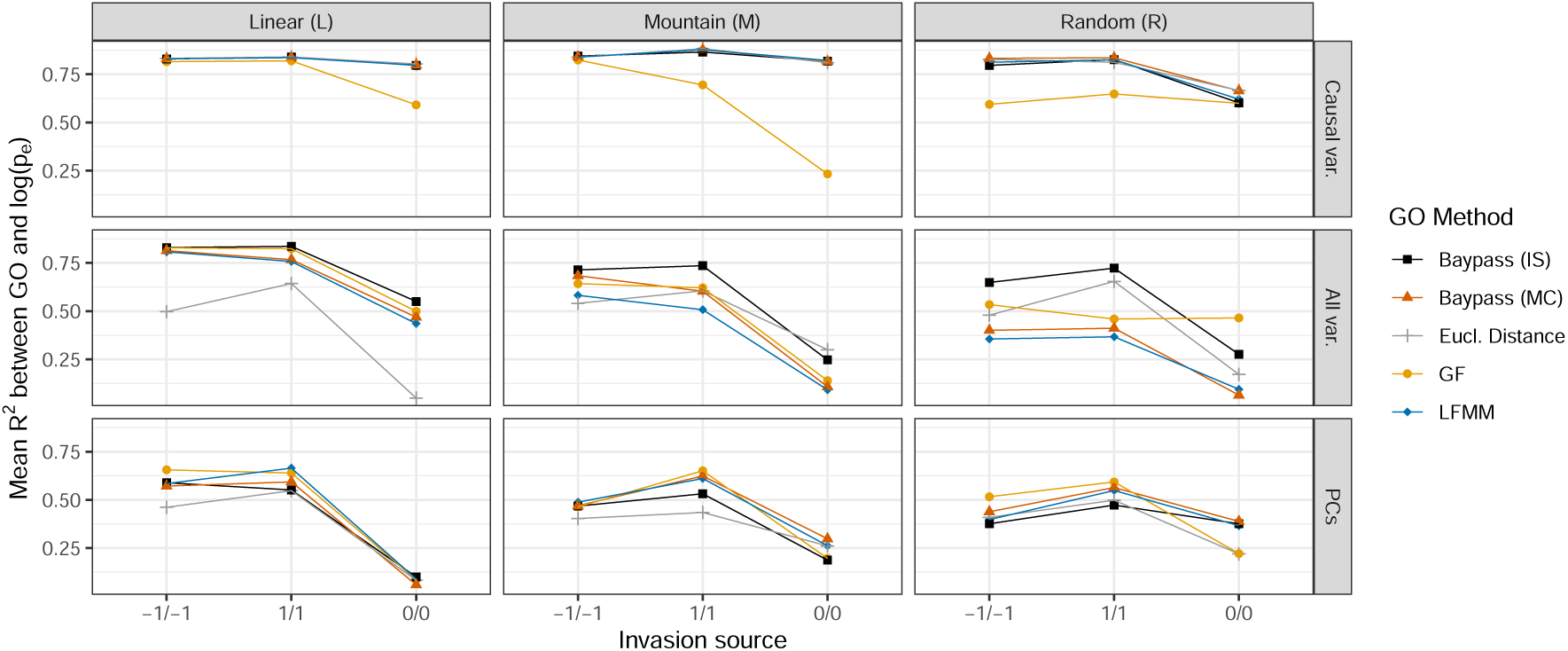
M**e**an **R**^2^ **values between GO and** log(*p_e_*) (for the low migration rate and 10 invading individuals) for the different native environment types (L on the left; M in the middle; and R on the left) as a function of the covariables included in the computation of GO (two causal variables on top; eight covariables including two causal and six confounding covariables on center; and five PCs at bottom). Each panel represents the mean R^2^ value over 90 observations for each of the 3 possible source population for invasion; specified on the x-axis; and over ten replicated simulation for the different GO estimators (GO_gf_, gGO_lfmm_, gGO_is_ and gGO_mc_, see the main text for details) alongside with Euclidean environmental distance. All SNPs were used for GO computation.

#### 3.1.1 Superior performances of gGO measures when computed on causal variables

The R^2^ values between log(*p_e_*) and gGO remained consistently high when using only causal variables to compute gGO, with R^2^ exceeding 0.75 in most cases. Overall, there was no noticeable difference between the performances of the 3 gGO estimators (gGO_lfmm_, gGO_mc_ and gGO_is_). In contrast, GO_gf_ exhibited less consistent performance than gGO and the Euclidean distance, with a R^2^ value of only 0.25 in the worst case. Notably, R^2^ values between log(*p_e_*) and gGO using causal variables were similar to those between log(*p_e_*) and Euclidean distance, as previously observed by e.g. Ĺaruson *et al*. (2022). Indeed, one main interest of using GO measure is to weight variables according to their genetic importance. In the ideal scenario where only causal variables are employed for its computation, GO is not anticipated to demonstrate better performance than the Euclidean distance, as there is no need to discern which variables are related to adaptation. Despite their unrealistic nature, these results demonstrate the existence of a relationship between GO and log(*p_e_*) under ideal conditions. Additionally, it is noteworthy that for the 0/0 source population, R^2^ values tended to decrease in comparison to the other two source populations. This reduction can be attributed to the fact that these populations, with environmental values equal to 0, occupy a mid-range position within the spectrum of possible environmental values. This positioning results in less extreme GO values as well as less variable establishment probability values, making the ranking more challenging.

#### 3.1.2 Robustness of GO measures to confounding covariables

When introducing additional confounding variables in the GO computation, differences between the different GO methods and the Euclidean distance became more apparent. R^2^ values for all methods remained relatively high in the L environment, exceeding 0.75 for the –1/-1 and 1/1 source populations, and notably outperformed the Euclidean distance. However, for the M environment, R^2^ values decreased for all GO methods, with the majority falling below 0.75. For the R environment, the results exhibited a more significant decline. gGO_lfmm_ and gGO_mc_ R^2^ values dropped to around 0.5 and decreased to less than 0.125 in the case of the 0/0 source population, while Euclidean distance yielded higher R^2^ values than these two gGO methods. Conversely, GO_gf_, exhibited higher R^2^ values compared to Euclidean distance for the 0/0 and –1/-1 source populations. Interestingly, the gGO_is_ generally outperformed all other methods and maintained R^2^ values close to 0.75 in all environments for the 1/1 and –1/-1 source populations (but see below).

#### 3.1.3 Using PCs reduce differences in method performances

When computing GO on PCs of (true and confounding) variables, the results showed similarities to those obtained using all variables, but the differences between methods were less pronounced and R^2^ values were lower. In the case of the L environment, the results were slightly less favorable than with all variables, ranging between 0.5 and 0.75 for all methods and the 1/1 and –1/-1 source populations, but dropping to less than 0.125 for the 0/0 source population. For the M and R environments, the reductions in R^2^ values were less pronounced, with most R^2^ values remaining relatively similar to those observed when using all variables. Moreover, employing PCs resulted in a narrower performance gap between gGO_is_ and other gGO methods, occasionally leading to gGO_is_ being outperformed by alternative methods. Conversely, when using PCs, Euclidean distance exhibited comparable or slightly worse performance compared to GO methods.

#### 3.1.4 Impact of covariables correlation on the gGO estimators

It might seem at first surprising that gGO_is_ performed equally (when considering the two causal covariables only) or even better (when considering all eight covariables) than gGO_mc_ and gGO_lfmm_ since the underlying GEA models are similar and previous work showed that the IS estimation of regression coefficients was suboptimal compared to MCMC based estimation (Gautier, 2015). However, the trend was less clear when considering PCs suggesting a possible negative impact of the correlation of covariables in the estimation of gGO_mc_ and gGO_lfmm_, both relying on a joint modeling of all the covariables while the IS estimation of regression coefficients (used to estimated gGO_is_) amounts to treat all covariables separately. We thus hypothesized that the suboptimal performance of gGO_mc_ and gGO_lfmm_ when additional confounding variables were introduced, might be related to a poorer estimation of regression coefficients of correlated covariables. To explore this, we conducted a comparative analysis between BayPass IS, BayPass MC, and LFMM. However, this time, we estimated regression coefficients independently for each variable (i.e., running BayPass MC and LFMM separately for each covariable). Results for a low migration and 10 invading individuals are shown in Figure 3. Univariate calculation of regression coefficients resulted in clear improvements for gGO_lfmm_ and gGO_mc_, aligning them closely with gGO_is_ in most instances, notably in the M and R environments. Although gGO_is_ mostly maintained slightly higher R^2^ values, particularly in the L environment, gGO_lfmm_ occasionally outperformed it, as seen in the M and R environments for 0/0 source population.

**Figure 3:**
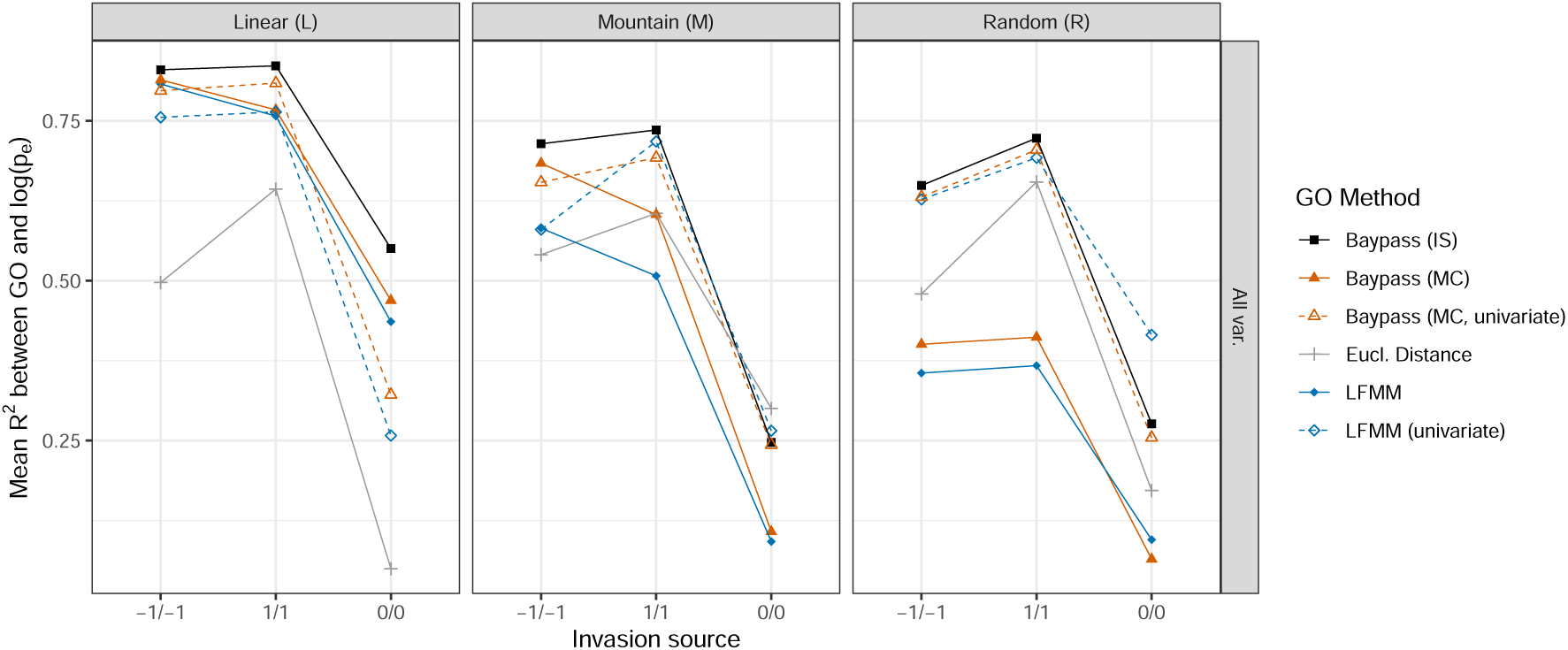
M**e**an **R**^2^ **values between GO and** log(*p_e_*)**, for the low migration rate and 10 invading individuals**, depending on the type of native environment (L, M or R). Each panel represents the mean R^2^ value (for each of the 3 possible source population for invasion) between log(*p_e_*) and GO obtained through five gGO computation methods, including gGO_lfmm_ and gGO_mc_ modified in order to treat variables independently (noted as “univariate”), alongside Euclidean distance. All SNPs and covariables were used for GO computation.

#### 3.1.5 Influence of SNPs pre-selection and simulation parameters

Overall, with the gGO method, the effect of SNP pre-selection on performance was not clear, as it did not consistently lead to better or worse R^2^ values. However, when using GO_gf_, a clear trend emerged where SNPs pre-selection consistently resulted in lower R^2^ values compared to cases where SNPs are not pre-selected (Figure S5 and S6). Similar to the SNPs pre-selection, the migration rate had a small impact on the results. Whether the migration rate was low (Figure 2) or high (Figure S7), the results remained overall quantitatively and qualitatively comparable. Indeed, for high migration rate, substantial R^2^ values were observed across all GO methods when exclusively employing causal variables, and a decline in R^2^ values occurred upon the addition of confounding variables. As for the number of invading individuals, while a linear relationship between log(EP) and GO is observed for 10 invading individuals, the same does not hold when the invading population size increases to 100 (see Figure S3 for an example). In the latter case, where EP values are consistently close to one, calculating R^2^ values becomes less meaningful. That is why the R^2^ results for the 100-individual scenario are not presented here. However, it is noteworthy that the results regarding the association between GO and fitness, as well as GO and population growth rate, exhibited comparable outcomes than the ones between GO and establishment probability (Figures S8 and S9), and that a robust relationship between GO and fitness or growth rate persisted even with 100 invading individuals, as depicted in Figure S10.

### 3.2 Interpreting gGO absolute value in terms of *f*_2_

To provide insights into the biological interpretation of gGO, we compared the accuracy of the different gGO estimators based on the theoretical expectations of Gain *et al*. (2023), who showed that the value of gGO between two locally adapted populations should be equal to the *f*_2_ measured at causal QTNs for these two populations.

Using all QTNs and the two causal environmental variables yielded estimated gGO closely related to *f*_2_ in the L environment, with small MAPE for both the gGO_lfmm_ and gGO_mc_ estimators across varying migration rates (Figure S11). For instance, for the highest migration rate, the gGO_lfmm_ estimator deviate from the true *f*_2_ by only 17%. Predictions were less accurate for the M environment, where gGO_lfmm_ MAPE value reached 64% in the case of low migration, and for the R environment with MAPE ranging from 69% to 81% (Figure S11). gGO_mc_ and gGO_lfmm_ exhibited very similar results, with gGO_mc_ appearing slightly more accurate in most cases.

In practice, calculating GO on all causal SNPs only is unrealistic notably because the driving co-variables are usually not all included in the analysis and even in this case, no GEA method could be expected to classify perfectly (i.e., with a decision criterion leading to a power of 1 and a no false discoveries) all the underlying associated SNPs. The results of the comparison between a more “realistic” gGO computed using both causal and neutral SNPs (with a MAF filter) and the *f*_2_ computed either with QTNs onlys or with both QTNs and neutral markers are presented in Figure 4. For improved readability and to reduce computational intensity associated with computing allele frequencies for all individuals and markers, we present results for only two environmental seeds and the L environment. Results for the M and R environments are shown in Figure S12. These “realistic” gGO calculations consistently overestimated the *f*_2_ calculated for QTNs alone, suggesting an imperfect estimation of the regression coefficients leading to wrongly consider some neutral QTNs as adaptive. MAPE values were overall quite high, indicating limited accuracy in predicting true adaptive *f*_2_ values. As already observed for relative establishment probabilities, absolute gGO values were the most accurate in the L environment and decreased in the M and R environments. On a more positive note, we observed that estimated gGOs were clearly lower than the *f*_2_ computed with both QTNs and neutral markers, implying that these categories of SNPs could be at least partly distinguished.

**Figure 4:**
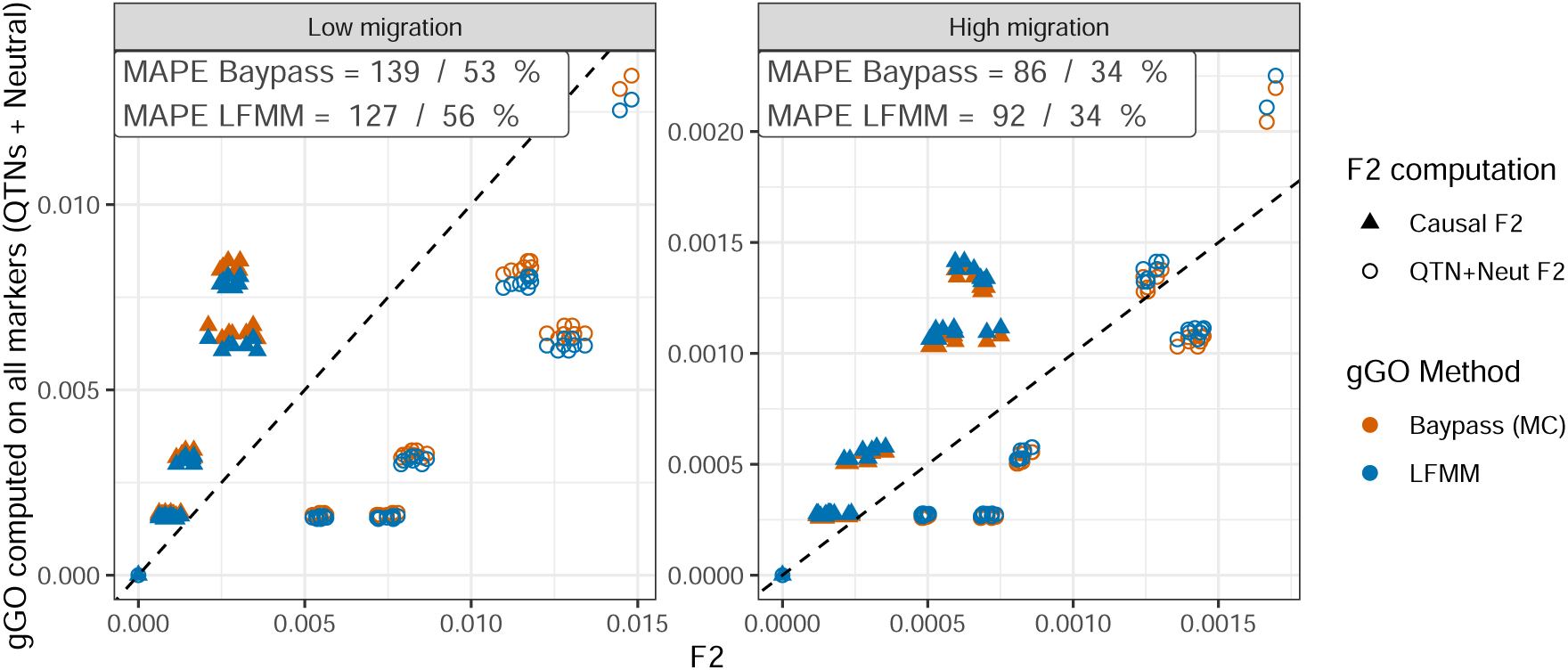
C**o**mparison **between gGO and** *f*_2_ **among QTNs (i.e. “ground truth” GO value) or among QTNs and neutral SNPs**. The estimated values with the two gGO estimators (gGO_lfmm_ and gGO_mc_) were obtained using QTNs and neutral markers (with MAF*>* 0.01) for the scenarios with Low (left panel) or High (right panel) migration within the native area under the L (linear) environment. The inset in each panel gives the two corresponding MAPE separated by a slash.

### 3.3 *B. tryoni* case study

GO computations conducted between the source population (population 1) and an extensive area in Oceania, employing PCs as predictor variables, demonstrated consistent outcomes across gGO_mc_, gGO_lfmm_ and GO_gf_ (Figure 5).

**Figure 5:**
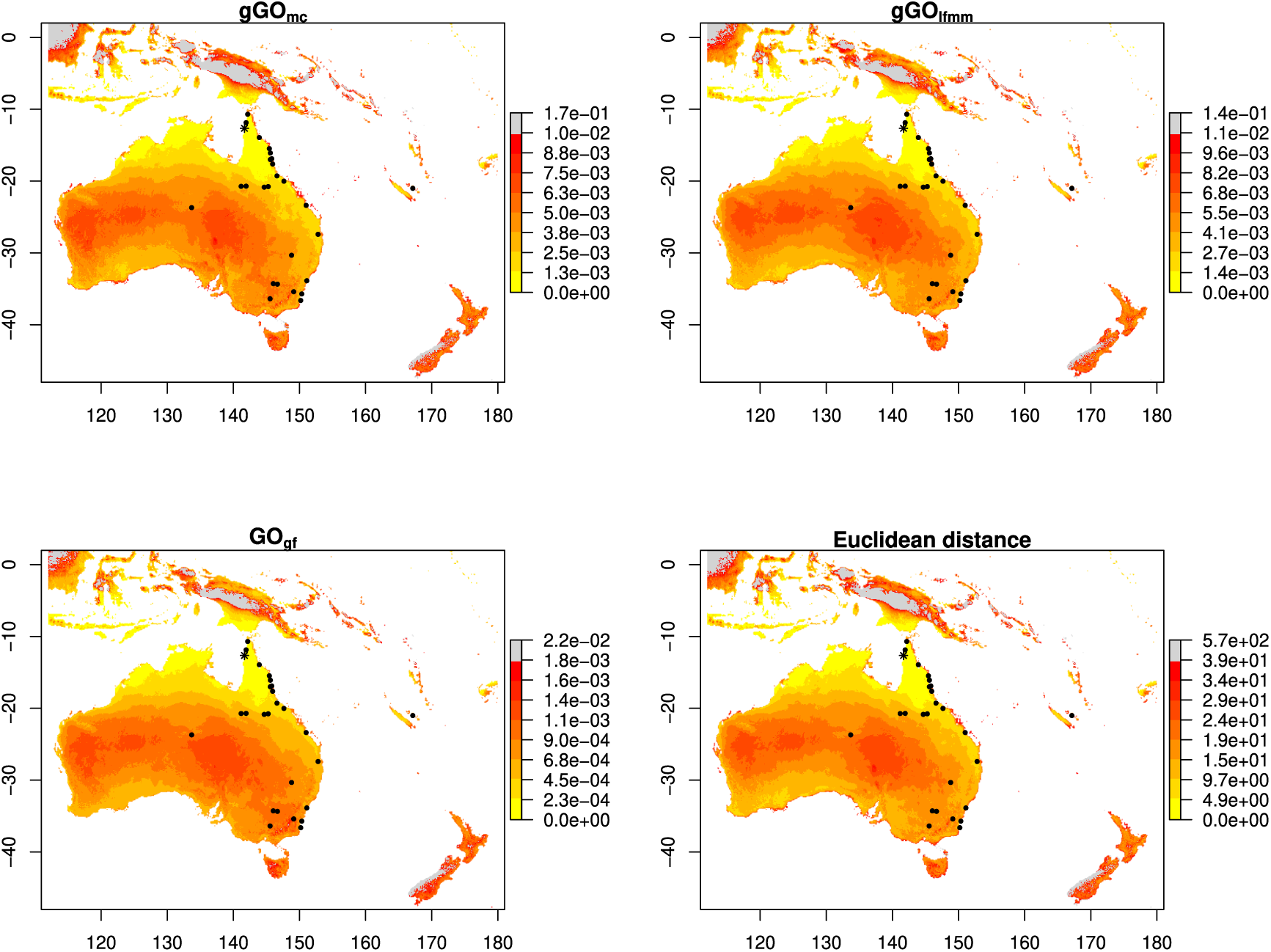
A**p**plication **of GO to *B. tryoni* populations.** GO was estimated between population 1 (“source” population, identified with a star) and a large area in Oceania, with gGO_mc_, gGO_lfmm_ and GO_gf_. Squared Euclidean distance to the source population is also displayed. Grey pixels represent outliers values, and black dots the studied populations.

Notably, all three methods identified outlying geographical zones in the North West area of New Zealand, the West part of Indonesia and in the central region of Papua New Guinea, while also identifying New Zealand and Tasmania as being some of the areas having the highest GO values. Similarly, all methods exhibited the lowest GO in the North of Australia, Southern Papua New Guinea, South of Western Australia, and Southern Indonesia islands. Islands situated to the east of Australia (comprising Loyalty and Fiji Islands) exhibited medium GO. Some of the highest GO values in mainland Australia concentrated in the central part of Western Australia and the Eastern region shared by Northern Australia, Queensland, and South Australia. These “visual” similarities are confirmed by a high correlation of all GO estimators values, particularly between gGO_lfmm_ and gGO_mc_ (Figure S14). Euclidean distance showed similar results to all GO methods, notably showing high correlation level with gGO_lfmm_ and gGO_mc_ (Fig. S14).

When using the six environmental covariables selected by Parvizi *et al*. (2023), results for multivariate methods closely resembled those obtained with PCs (Figure S15). Disparities between “univariate” and “multivariate” methodologies were also observed in agreement with the simulation study. More precisely, results obtained using univariate methods (gGO_lfmm_ “univariate” and gGO_is_) differed from those obtained using PCs, likely due to correlation between variables. Among multivariate methods, gGO_mc_ and gGO_lfmm_ were again very similar and overall identified the same low/high GO regions as those identified with PCs. GO_gf_ differed from these two approaches, for example not showing high GO regions in the central part of Western Australia and the Eastern region shared by Northern Australia, Queensland, and South Australia. Again, Euclidean distances gave comparable results to gGO_lfmm_ and gGO_mc_ but displayed lower correlations compared to those observed when using PCs (Figure S16).

## 4 Discussion

The purpose of this study was to conduct a comprehensive evaluation of GO measures, specifically focusing on their performances within the framework of biological invasions. The primary objective was to determine whether GO measures could effectively predict establishment probabilities through a simulation study. Furthermore, the application of GO measures to the biological invasion of *B. tryoni* served to illustrate their practical utility in this context. Finally, our study also sought to provide insights into the feasibility of interpreting the absolute values of gGO measures, while introducing methodological innovations through the computation of gGO with BayPass software and the optimization of Gradient Forest R package.

### 4.1 Predicting relative establishment probabilities using GO

In general, at least one GO measure outperformed Euclidean distance in predicting EP, except when predicting EP using causal variables only, where all approaches performed similarly. These results were expected, as the unrealistic scenario where causal variables are known suppresses one of the main advantages of GO: weighting variables according to their genetic importance. The better performance of GO methods over Euclidean Distance when confounding variables are added underlines the effectiveness of using genetic information to improve the prediction of establishment probabilities, providing insights into the genetic make-up of the populations studied.

While our simulations indicate overall good performances of GO in predicting EP, these performances differed between native environments. The L environment, defined by gradual transitions between populations’ environmental optima, tends to generate clearer relationships between allele frequencies and the environment. Conversely, the M environment contains geographically distant populations with similar environments (e.g., populations 1, 5, 21, and 25, Fig. 1), and the R environment juxtaposes dissimilar environments. This complexity may lead to populations with different genetic composition under similar environmental conditions due to low migration, or with similar genetic composition under different environments due to high migration, likely resulting in slightly poorer GO performances. Moreover, our simulations were characterized by high polygenicity and genetic redundancy, where numerous potentially pleiotropic QTNs were segregating and multiple combinations of these QTNs could lead to the same trait values. The interplay between this high polygenicity and the complex patterns of local adaptation in the M and R environments likely explains the poorer performances of GO methods in these scenarios. These results are in line with those of Lotterhos (2023), illustrating that high polygeny, genotypic redundancy and pleiotropy can result in non-monotic patterns between allele frequencies and environmental variables. Such patterns challenge GEA methods, which rely on the assumption of clinal patterns between allele frequencies and environmental variables.

Overall, gGO methods outperformed GF in predicting EP, especially when each variable was considered individually in gGO calculations. This is in line with the findings of Gain *et al*. (2023), who focused on the relationship between GO and fitness. In the ideal case with only causal variables, GO_gf_ often showed lower performance than Euclidean distance, indicating limited ability to decipher the allele frequency-environment relationship. However, the performance gap between gGO and GF decreased when confounding variables or PCs were included in GO calculations. In scenarios where non-linear relationships exist between allele frequencies and environment, GO_gf_ may in theory out-perform gGO due to its ability to accommodate such relationships. While this could not be clearly observed in the more complex M and R environments of our simulations, we note that GF is a machine learning based method whose performance is likely more dependent on the amount of data; thus, we cannot rule out that our dataset of 25 populations may have been insufficient to make accurate predictions with this approach. As underlined by Gain *et al*. (2023), linear model may also achieve a better bias-variance trade off than non-linear machine learning model.

Among gGO methods, gGO_mc_ and gGO_lfmm_ yielded similar results, which was expected as these methods are based on the same principle. In practice, although LFMM is more computationally efficient, the Bayesian hierarchical framework underlying BayPass allows to accommodate and properly account for the specificities of non-standard (e.g., Pool-Seq data) or heterogeneous data sets (Camus et al., in prep), thereby opening new ways to apply gGO in some biological contexts. Likewise, promising directions would be to directly incorporate estimation of the matrix ***B*** of the SNP environmental effects (regression coefficients) in the model to allowing accounting for correlation among covariables (if not using PCs), and to provide estimates of uncertainty (e.g., credibility interval) associated with the estimated GO.

In the presence of confounding variables, a noteworthy finding was that the prediction power of gGO methods was reduced only if regression coefficients were estimated jointly (i.e. the multivariate approach), but not if these coefficients were estimated independently for each variable (i.e. the univariate approach). The relatively stable performance of univariate methods can be related to the theoretical result of Gain *et al*. (2023), stating that a gGO computed from linear combinations of causal (and potentially non-causal) variables should be equivalent to the gGO computed with causal predictors only. Indeed, the variance-covariance matrix of regression coefficients that is used in gGO allows mitigating the redundancy of information resulting from the inclusion of both causal and correlated variables, while removing the noise arising from fake uncorrelated variables. However, a strong assumption underlying these expectations is that regression coefficients of observed variables are correctly estimated. This might explain the lower performance of multivariate approaches in our simulations, because the joint estimation of regression coefficients from a set of correlated variables is typically more challenging.

While these results suggest to always favor univariate methods to ensure accurate coefficient estimates, note that their prediction accuracy is certainly boosted by the inclusion of true causal variables in gGO computations, which is very unlikely in real-life studies. In comparison, simulation results based on PCs of environmental variables might better reflect the practical performance of GO methods. Prediction accuracy decreased for all methods in this case, and the differences between methods were also reduced. This result was expected since PCs potentially cause a loss of information about causal variables but avoid estimation issues by removing correlations between variables. Nevertheless, PC based GOs maintained an R^2^ value close to 0.5 (for –1/-1 and 1/1 source populations), which can be considered an acceptable predictive performance.

We therefore suggest caution when using multivariate gGO methods, and recommend either the use of PCs to suppress variable correlations or the use of univariate gGO for untransformed variables, even though the behavior of this latter approach with real data featuring many highly correlated variables and possibly no causal ones remains uncertain. A limitation associated with the use of PCs lies in the interpretation of variable importance, a facet of interest in GO approaches, because GEA models then report the importance of PCs, which have no clear biological interpretation. To overcome this issue, note that obtaining the importance of original variables from those of the PCs is actually straightforward (Supplementary Note 3). Finally, the impact of pre-selecting markers (e.g., based on XtX*^*^*) on gGO performance was found to be negligible, while it could compromise GF’s performance, affirming previous findings Gain *et al*. (2023). Therefore, we propose that pre-selecting markers is not an obligatory step for achieving a robust GO interpretation.

Despite the overall good performances of GO methods to predict EP, the scenarios considering the invasion of individuals originating from the 0/0 source populations highlighted some more conceptual limitations of GO-based EP prediction. Indeed, EP prediction was more difficult for 0/0 source populations, because their mid-range environmental values imply a relatively low adaptive challenge whatever the invaded environment. For instance, in the L environment with a low migration rate and 10 invading individuals, EP was approximately equal to one third in the most extremes 1/1 or –1/-1 environments (Figure S17). Considering causal variables only, a strong correlation between GO and EP was found despite the quite similar values associated to the different invaded environments (Figure S17). However, the additional noise resulting from the inclusion of confounding variables was sufficient to affect the ranking of GO values among invaded environments, which lead to a strong decrease of R^2^ values. This illustrates the problem of the lack of interpretability of absolute GO values. In such instances, a GO analysis should ideally conclude that *all* environments present a high invasion risk, not only that 0/1 environments are more at risk than 1/1 environments (for instance).

In other words, interpreting absolute GO values would be crucial to determine whether the variations of GO computed in distinct environments imply distinct or similar challenges for adaptation.

Unfortunately, our attempt to evaluate the interpretability of absolute gGO values outlined the difficulty of this task, with gGO values often diverging from their expected *f*_2_ values. This may be due to inaccurate estimation of regression coefficients, stemming from a variety of factors such as the small number of observed population, the existence of non-monotonic clines between allele frequencies and covariates or imperfect correction of population structure. Deviations from the conditions where gGO is expected to equal *f*_2_ (namely the infinitesimal model) may also contribute to this imperfect match. Nonetheless, our findings are promising as they reveal a strong proximity between gGO and *f*_2_ within the linear environment, under ideal conditions where only QTNs were considered. Moreover, a strong correlation between gGO and *f*_2_ persists across all environments, even in more realistic conditions where gGO was computed from all SNPs. In these conditions, gGO also exhibited some expected behavior as it overestimated the QTNs’ *f*_2_ (likely due to erroneously attributing weight to neutral SNPs) while remaining below the overall *f*_2_ (i.e., computed based on both QTNs and neutral SNPs) thus correctly excluding (or down-weighting) most neutral SNPs.

### 4.2 Towards a more comprehensive modelling of population fitness

Our simulation framework demonstrated a strong correlation between GO and EP across various environment types and migration rates, even when considering confounding variables. This sug-gests, among others, that GO is resilient in scenarios with moderate population differentiation and imperfect adaptation, such as those with high migration. However, several open questions need to be addressed before applying GO to predict EP beyond idealized simulation frameworks.

All GO methods assume that populations adapt to new environments through pre-existing variants, so their ability to predict fitness and thus EP is expected to decrease if adaptation actually proceeds, at least partly, from de novo mutations. However, the simulations conducted in this study do not allow quantifying this effect, because their design implies that adaptation is mainly driven by standing variation. Indeed, the polygenic traits’ architecture and the environment heterogeneity create high levels of standing genetic variation in the native area (Yeaman, 2022; Höllinger *et al*., 2019), facilitating rapid adaptation in invaded environments (Jain and Stephan, 2017). In addition, the relatively low mutation rates and the small founding population sizes make adaptation through *de novo* mutations very unlikely in the invaded area, at least in the short evolution time considered here. While a growing body of literature supports the idea that rapid adaptation to environmental change often results from standing genetic variation (Barrett and Schluter, 2008; Bitter *et al*., 2019; Chaturvedi *et al*., 2021), and deems it important for invasive species to adapt to invaded areas (Prentis *et al*., 2008; Bock *et al*., 2015), at least one reported case indicates adaptation through potential new mutations during colonization (Exposito-Alonso *et al*., 2018). Additionally, some adaptations to traits relevant to invasion biology, such as insecticide resistance in crop pests, are thought to result from the interplay between standing variation and *de novo* mutations (Hawkins *et al*., 2019). This emphasizes the importance of considering these processes when using GO methods. Additionally, when relying on GO to anticipate biological invasions, it is implicitly assumed that population pre-adaptation plays a significant role in the successful establishment in a new environment. However, scenarios with 100 invading individuals illustrate that it might not always be the case: while GO maintained a strong correlation with fitness (Fig. S10), accurate prediction of EP became challenging due to the substantial number of invading individuals buffering the adaptive challenges presented by the new environment. While meta-analyses have shown that invasive species often maintain their ecological niche in the invaded area (Bomford *et al*., 2009; Liu *et al*., 2020; Aravind *et al*., 2022), supporting the hypothesis that pre-adaptation plays a significant role in invasion success, successful population establishment can be influenced by various factors, including propagule pressure (Simberloff, 2009; Wittmann *et al*., 2014), hybridization/admixture (Rius and Darling, 2014; Barker *et al*., 2019) and epigenetic processes (Mounger *et al*., 2021; Marin *et al*., 2020). While our simulations partially explored the effects of propagule pressure by varying the number of invading individuals, the influence of successive introductions was not examined. Incorporating some of the above factors into simulations could refine the conditions under which GO can effectively predict EP in more realistic applications.

Our simulations also did not explicitly include recessive deleterious mutations, since QTNs could be either deleterious or beneficial depending on the environment and their genetic background always affects the phenotype. This hinders our ability to study the influence of genetic load on invasion success. However, biological invasions are generally characterized by initial bottlenecks favoring drift and potential inbreeding, which can reduce population fitness due to the fixation and/or expression of strongly deleterious mutations. Nevertheless, some studies have shown evidence of genetic load purging in invasive species (Facon *et al*., 2011; Tayeh *et al*., 2013; Mullarkey *et al*., 2013; Marchini *et al*., 2016), and gene flow and/or admixture can also mask genetic load (Whiteley *et al*., 2015). Furthermore, cases exist where populations, despite showing evidence of high genetic load without a clear purging signal, have successfully established and persisted in new environments (Gautier *et al*., 2023; Zayed *et al*., 2007). Demographic parameters and stochastic factors can influence genetic load and therefore population persistence in contrasting ways (reviewed in Robinson *et al*. (2023) and Bouzat (2010)). Further work is thus needed to explore the nuanced dynamics of invasion success in the presence of genetic load, as GO alone does not account for its effects.

### 4.3 Insights from the *B. tryoni* case study

Despite the limits mentioned above, practical use of GO in the case of *B. tryoni* appeared insightful. The results obtained with GO, especially with gGO_lfmm_ and gGO_mc_, are consistent with the existing knowledge of the species’ establishment. Consistently, the lowest GO values were obtained within the native range of *B. tryoni*, this area being not expected to pose any adaptive challenge for the chosen reference population. More interestingly, the medium GO estimated in the Loyalty Islands aligns with the early stages of expansion, as the first documented records in these islands date back to around 1969 (Popa-Báez *et al*., 2020). The higher, but still relatively low, GO values observed around population 26, populations 16 and 17, and the east coast of Australia, are consistent with a later establishment—circa 1987 for population 26 (Cameron, 2006) and approximately 1994 for populations 16 and 17, as well as the East Coast of Australia (Popa-Báez *et al*., 2020; Osborne *et al*., 1997).

The lower GO in areas not yet colonized (southern Papua New Guinea, southern Western Australia, and southern Indonesian islands) suggest that these regions may be vulnerable to establishment of individuals from the native area (population 1). Given its geographical proximity to the native range of *B. tryoni*, Papua New Guinea is at a higher risk of invasion. Regions with high GO in mainland Australia (for gGO_mc_ and gGO_lfmm_) coincide with areas where *B. tryoni* is not estab-lished. Nevertheless, in Western Australia, a few incursions have been documented, but eradication measures are promptly initiated upon the detection of more than five individuals, likely preventing the establishment of any population (Dominiak and Mapson, 2017). More broadly, it should be kept in mind that biocontrol strategies, which are frequently implemented (Maelzer *et al*., 2004), might have substantially impeded the establishment of *B. tryoni*, whether populations were pre-adapted or not.

High GO regions beyond mainland Australia also align with our knowledge of *B. tryoni*, as none of these areas exhibit established populations. However, Popa-Báez *et al*. (2021) have demonstrated that occasional incursions into New Zealand and Tasmania likely originated from New Caledonia or the east coast of Australia. Based on GO values, we could infer that the invasion risk stemming from Northern *B. tryoni* in these areas is low, but it is important to note that our analysis did not assess the invasion risk from other populations.

It is also important to highlight that, in this specific case study, incorporating genetic information to predict *B. tryoni* invasion risk only marginally affected the conclusions that would have been drawn by simply considering Euclidean distance between environmental covariables. This may reflect a limited genetic basis to adaptation in this species and/or a lack of genomic (number of SNPs) and environmental (available covariates) information captured by the specific dataset used to evaluate GO. Besides, employing PCs may lead to a loss of information regarding the association between genomic data and covariates, especially in real-life scenarios where causal variables are lacking. This could explain the higher correlation levels observed between GO methods and Euclidean distance that we observed when using PCs, compared to the less pronounced correlations when considering the set of ascertained untransformed covariates.

More generally, this case study effectively demonstrates a relatively straightforward but practical application of GO in a real biological invasion context.

### 4.4 General conclusions

Our study confirms the relevance of GO to predict invasion success and provides several methodolog-ical tools and advice to enhance the performance of this approach. Regarding empirical application, we illustrate how GO measures can be utilized to provide recommendations for invasion risk. Further theoretical research is needed to determine the impact of several key factors of invasion success, such as propagule pressure and genetic load, on the accuracy of predicting establishment probabilities with GO. Additional research on other species and how to integrate other genomic tools or SDM to provide a more comprehensive evaluation of invasion risk will also be necessary in the future.

## 5 Data Accessibility and Benefit-Sharing Section

The *B. tryoni* data have already been published, and have permissions appropriate for fully public release. The code necessary to reproduce the simulations, compute simulations GO, and the code to use the optimized version of Gradient Forest package is available at https://forgemia.inra. fr/simon.boitard/popgenomicprediction.

## Supporting information

Supplementary informations

## Acknowledgments

We wish to thank Olivier Fraņcois, Arnaud Estoup, Bénédicte Rhoné, Renaud Vitalis and Jöelle Ronfort for helpful insights and discussion. Louise Camus was funded by the Occitanie Region (France), and the INRAE Scientific Department SPE.

