## Supplementary informations for "Predicting species invasiveness with genomic data: is Genomic Offset related to establishment probability?"

### A Supplementary Figures

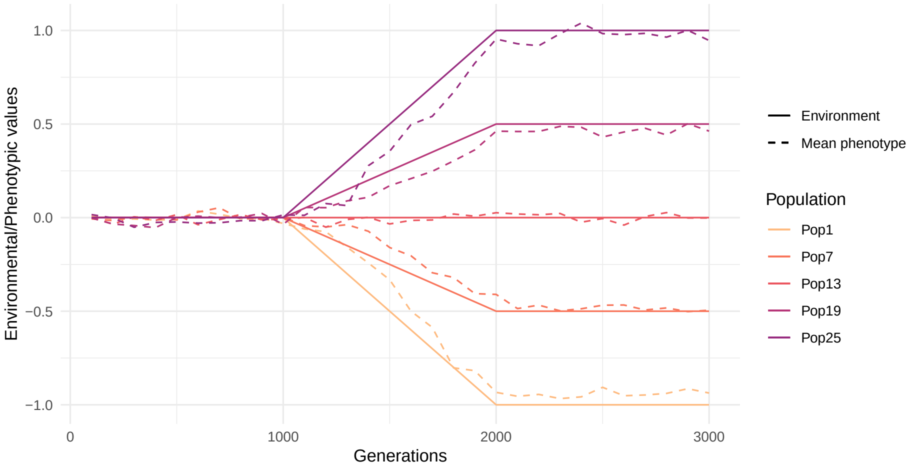


Figure S1: **Evidence of local adaptation in simulated native areas.** Evolution of environmental optimum 1 and mean realized phenotype along generations, for 5 populations of the native area simulations. A single random replicate is considered, for the scenario with low migration and environment type L. Populations indices are given in Figure 1.

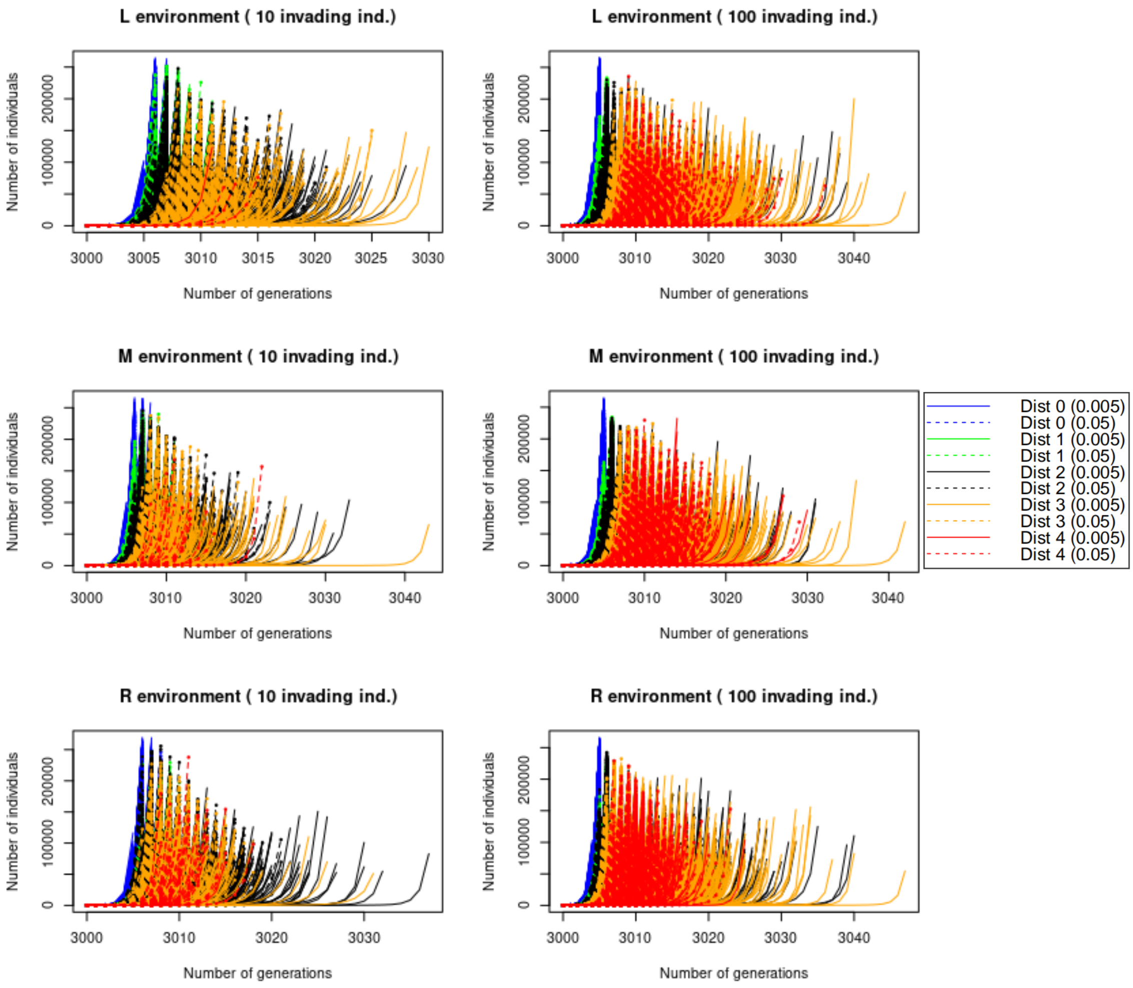


Figure S2: **Population size in the invaded area as a function of time (in generations)**, for the -1/-1 source population, across different environment types (L on the top, M in the middle and R in the bottom) and different numbers of invading individuals (10 on the right and 100 on the left). Populations always reach 50 000 individuals way before 100 simulated generations, and never face extinction after reaching this value. Colors correspond to the absolute environmental distance between the -1/-1 source population and the invaded environments (example : "Dist 2" includes the invaded environments at a distance of 2, either 0/0, 1/0 or 0/1). Solid lines represent the evolution of the number of individuals for low migration rate, and dashed lines for high migration rate.

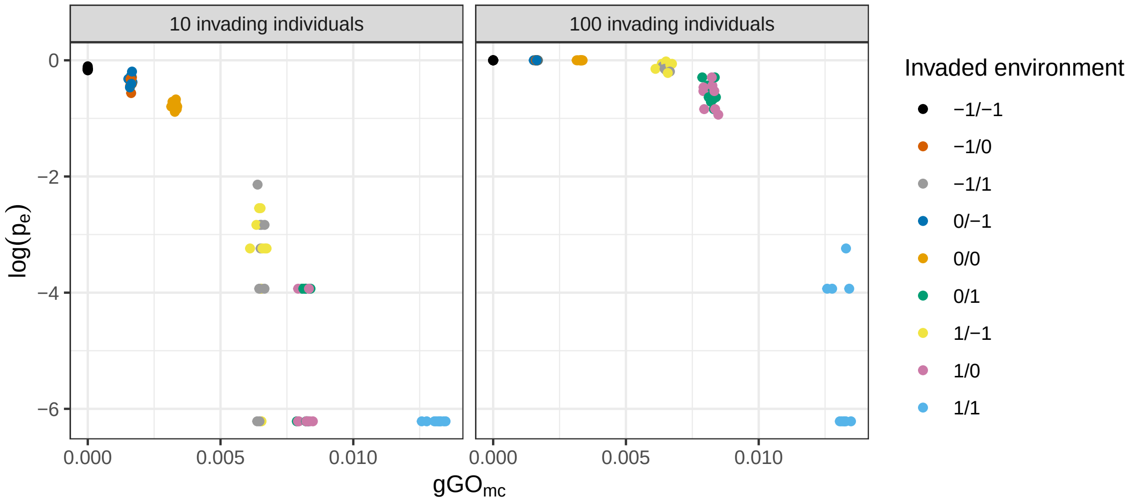


Figure S3: **Evaluating the correlation between GO (here $\mathbf{gGO_{mc}}$) and $\mathbf{log(\boldsymbol{p_e})}$.** On each panel, $\log(p_e)$ is plotted as a function of $gGO_{mc}$ for 90 observations (10 repetitions of the native environment scenario × 9 possible invaded environments), which are combined to compute one $R^2$ value. The left panel presents results for 10 invading individuals, and the right one for 100 invading individuals. In this example, the source population for the invasion is of -1/-1 type, migration rate is low, the native environment type is L, and only causal variables were used to compute $gGO_{mc}$.

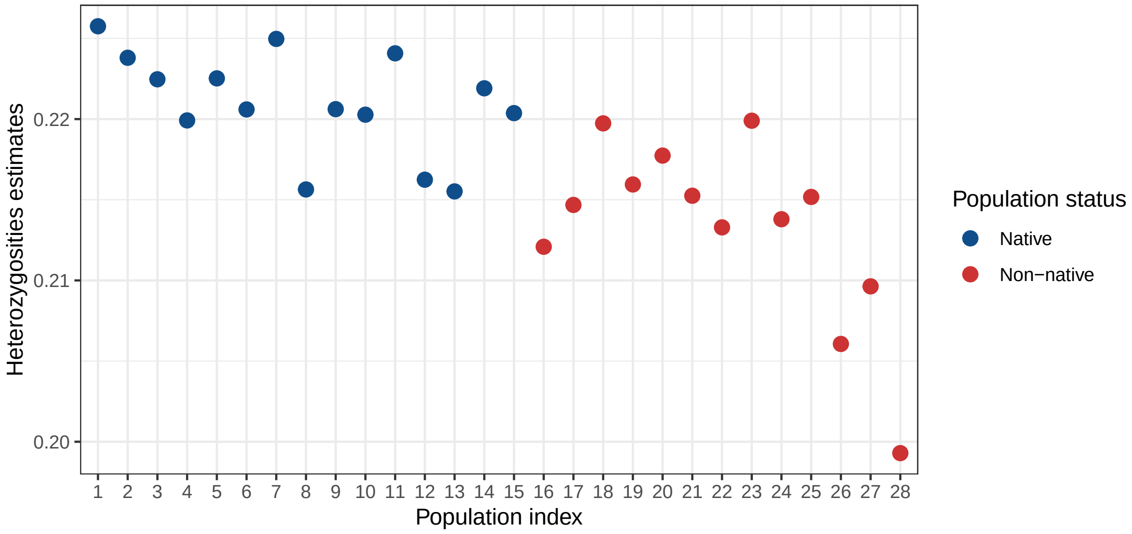


Figure S4: **Heterozygosities estimates computed with *poolfstat* for the 28 *B. tryoni* populations.**

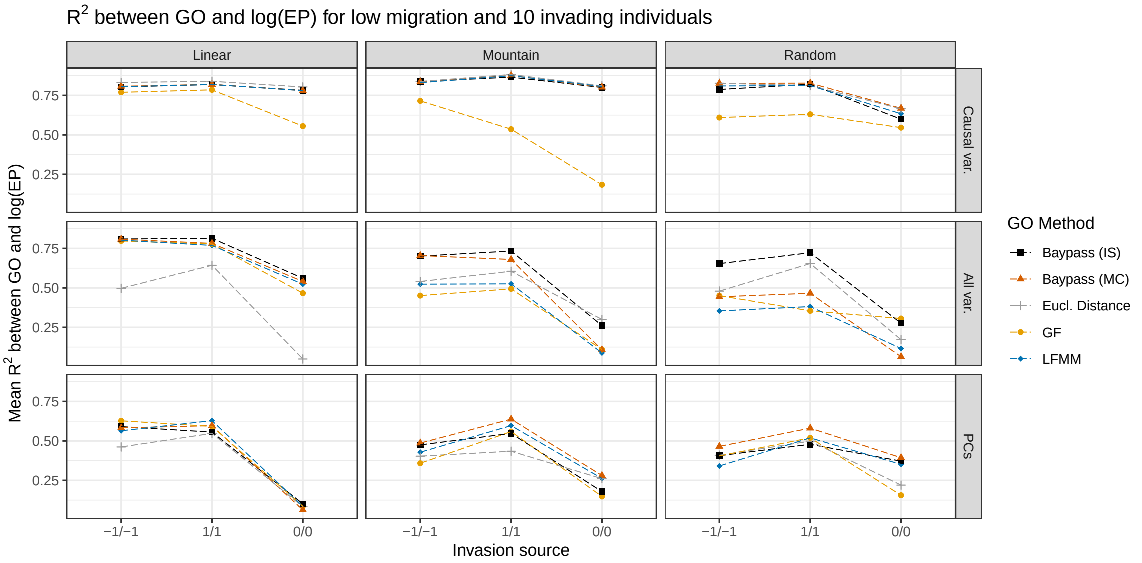


Figure S5: **Mean $R^2$ values between GO computed on top 10% XtX* QTNs and $\log(p_e)$ (for the low migration rate and 10 invading individuals) across the different native environment types (L on the left; M in the middle; and R on the left).** The results are presented as a function of the covariables included in the computation of GO (two causal variables on top; eight covariables including two causal and six confounding covariables on center; and five PCs at bottom). Each panel represents the mean $R^2$ value over 90 observations for each of the 3 possible source population for invasion; specified on the x-axis; and over ten replicated simulation for the different GO estimators ($GO_{gf}$, $gGO_{lfmm}$, $gGO_{is}$) alongside with Euclidean environmental distance.

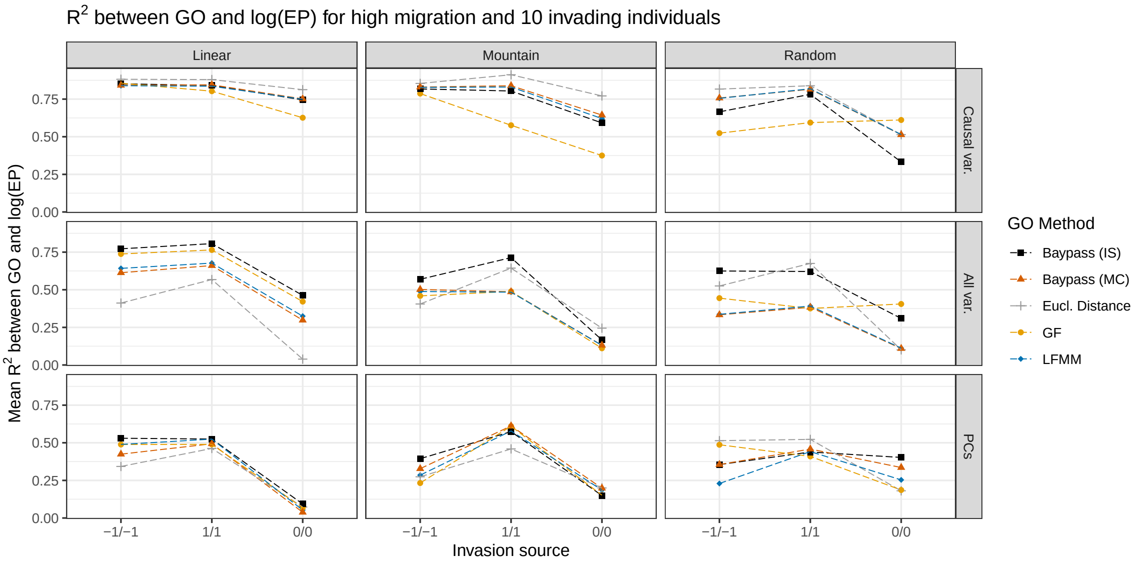


Figure S6: **Mean $R^2$ values between GO computed on top 10% XtX* QTNs and $\log(p_e)$, for high migration and 10 invading individuals.** Results are presented for different environment types (columns) and covariables included in the computation of GO (lines). Each panel represents the mean $R^2$ value over 90 observations for each of the 3 possible source population for invasion; specified on the x-axis; and over ten replicated simulation for the different GO estimators ($GO_{gf}$, $gGO_{lfmm}$, $gGO_{is}$ and $gGO_{mc}$, see the main text for details) alongside with Euclidean environmental distance.

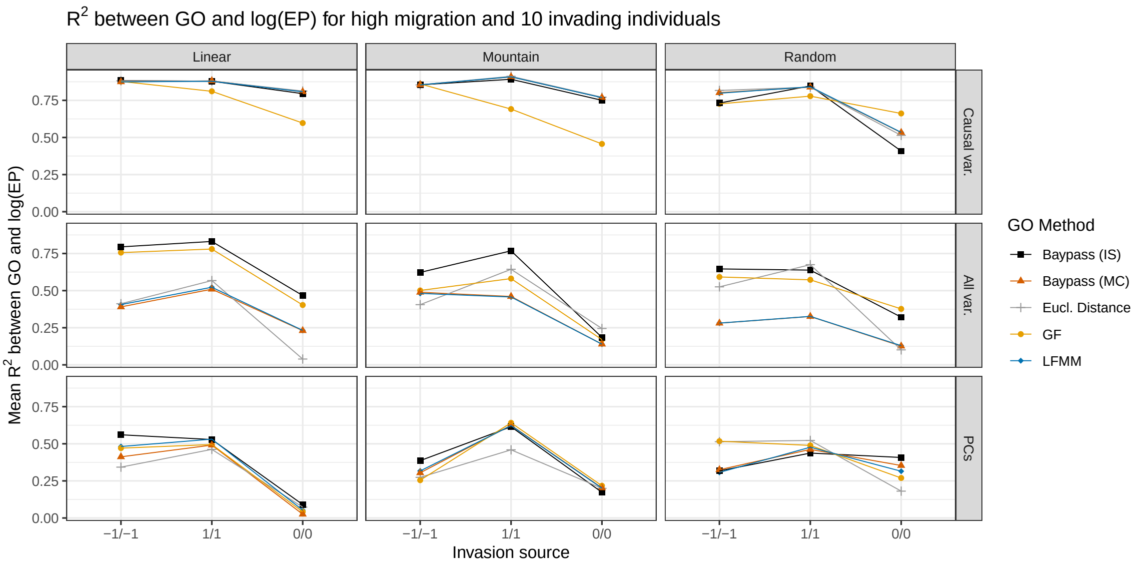


Figure S7: **Mean $R^2$ values between GO computed with all QTNs and $\log(p_e)$, for high migration and 10 invading individuals.** See Figure S6 for more details.

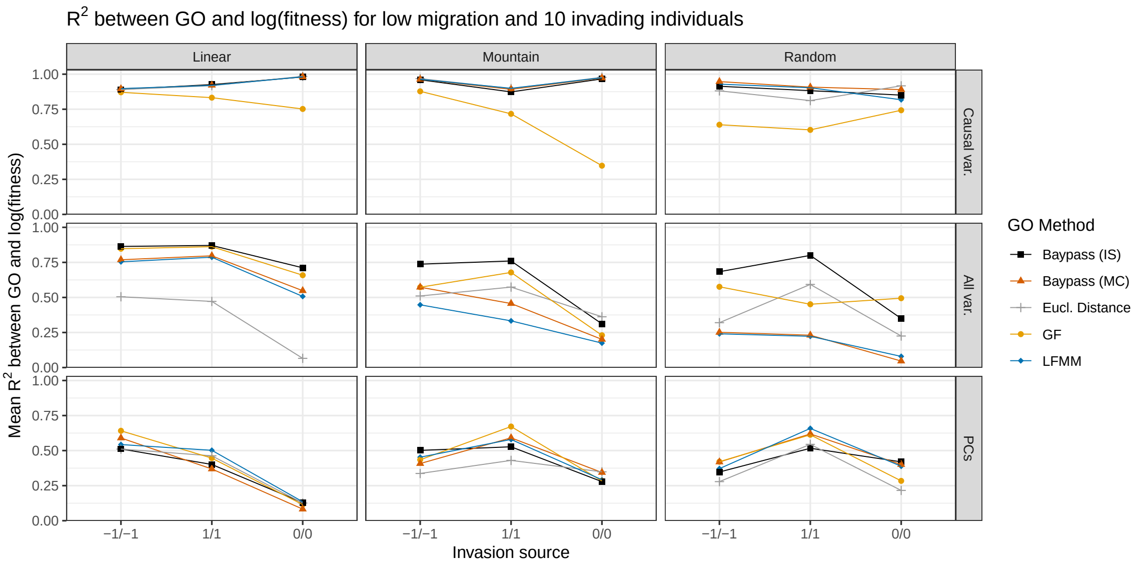


Figure S8: **Mean $R^2$ values between GO computed with all QTNs and log($fitness$), for low migration and 10 invading individuals.** See Figure S6 for more details.

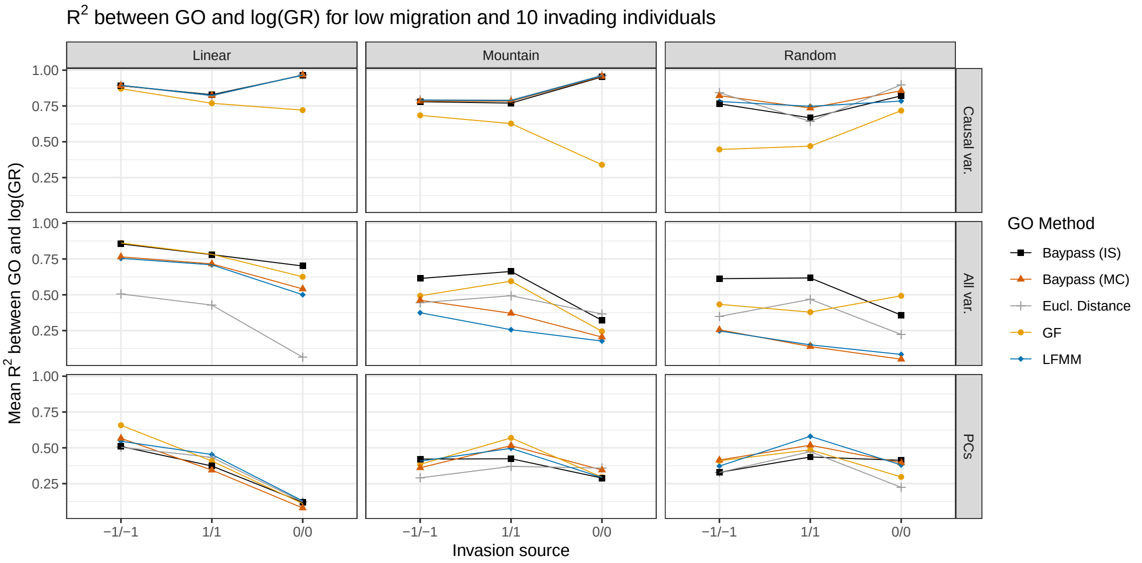


Figure S9: **Mean $R^2$ values between GO computed on all QTNs and log($growthrate$), for high migration and 10 invading individuals.** See Figure S6 for more details.

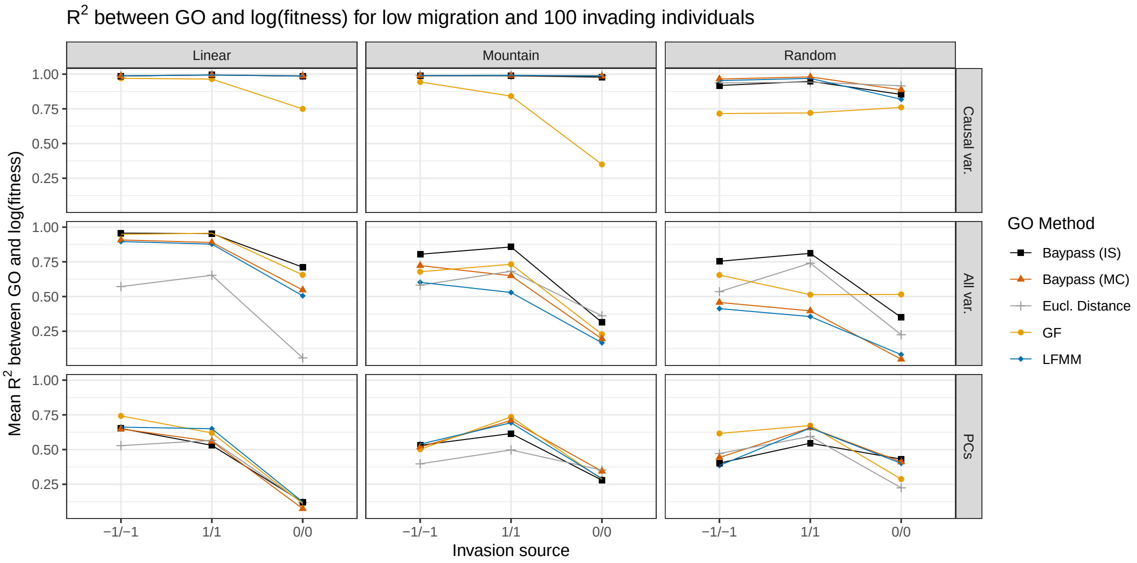


Figure S10: **Mean $R^2$ values between GO computed on all QTNs and log(*fitness*), for low migration and 100 invading individuals.** See Figure S6 for more details.

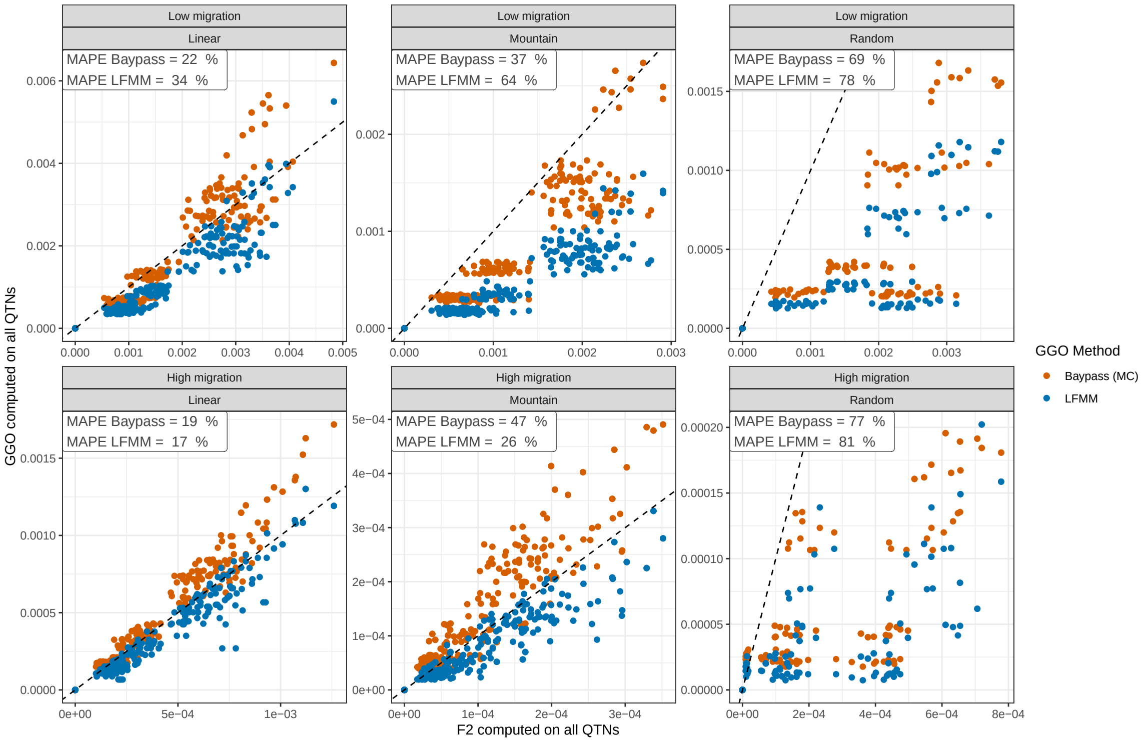


Figure S11: **Comparison of $\mathbf{gGO_{mc}}$ and $\mathbf{gGO_{lfmm}}$ with $f_2$ statistics**, for different native environment types (columns) and migration rates (lines). All causal QTNs were used for gGO and $f_2$ computation, without applying a MAF filter and excluding neutral markers. MAPE is indicated for each method and each type of scenario. The R environment dataset has fewer data points due to the absence of certain environmental value combinations in the native grid, resulting from the random selection of these values.

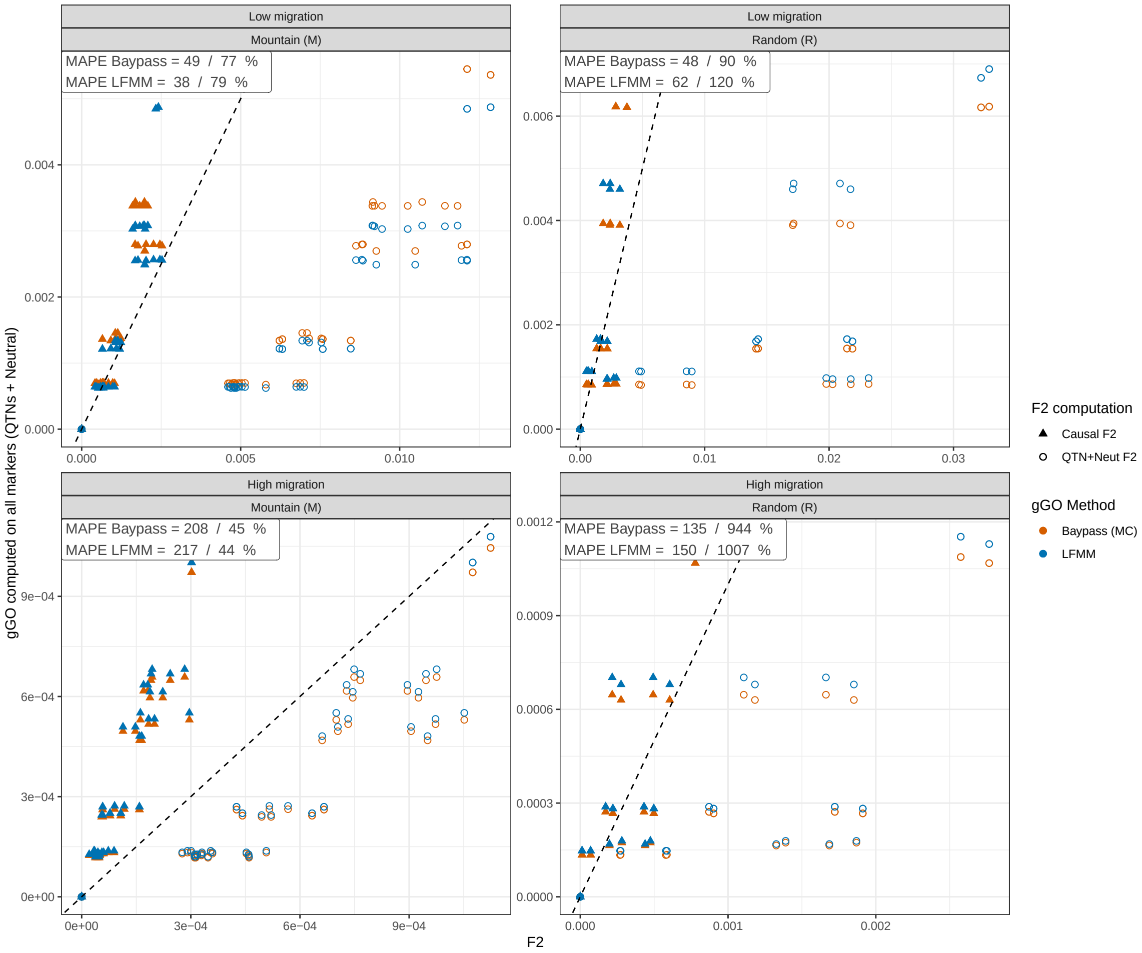


Figure S12: **Comparison of gGO computed using BayPass (MC) or LFMM with $f_2$ statistics derived from either QTNs alone or QTNs and Neutral markers (MAF-filtered), for M and R environments**. gGO computations involved QTNs and Neutral markers (MAF filtered). MAPE is indicated for each method and each type of scenario. The R environment dataset has fewer data points due to the absence of certain environmental value combinations in the native grid, resulting from the random selection of these values.

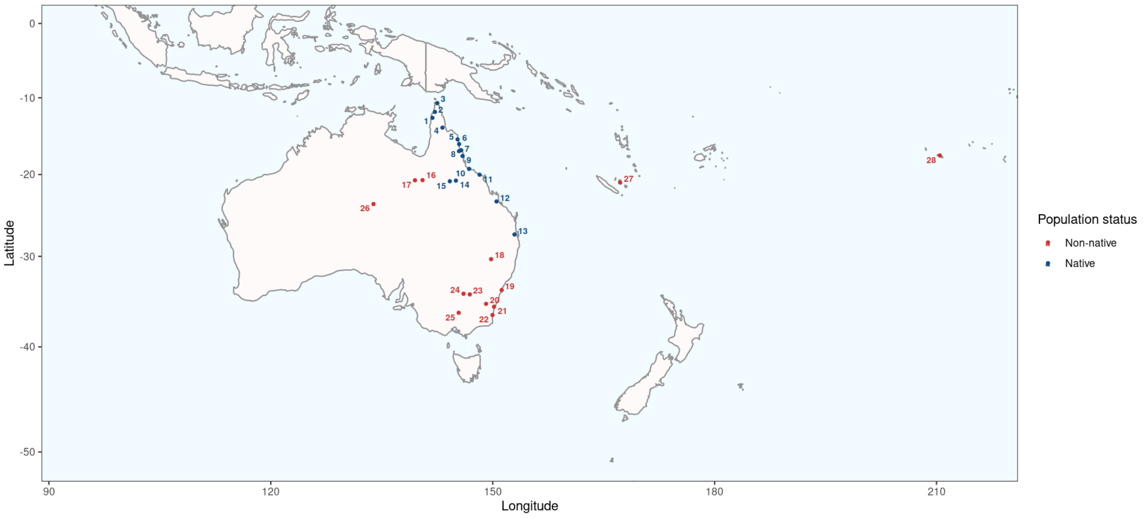


Figure S13: **Geographical Distribution of 28 *B. tryoni* populations studied by Popa-Báez *et al.* (2020)**. Populations 1 to 15 (in blue) are native populations, while populations 16 to 28 (in red) established outside the species' native range at different time points during the second half of the 20th century.

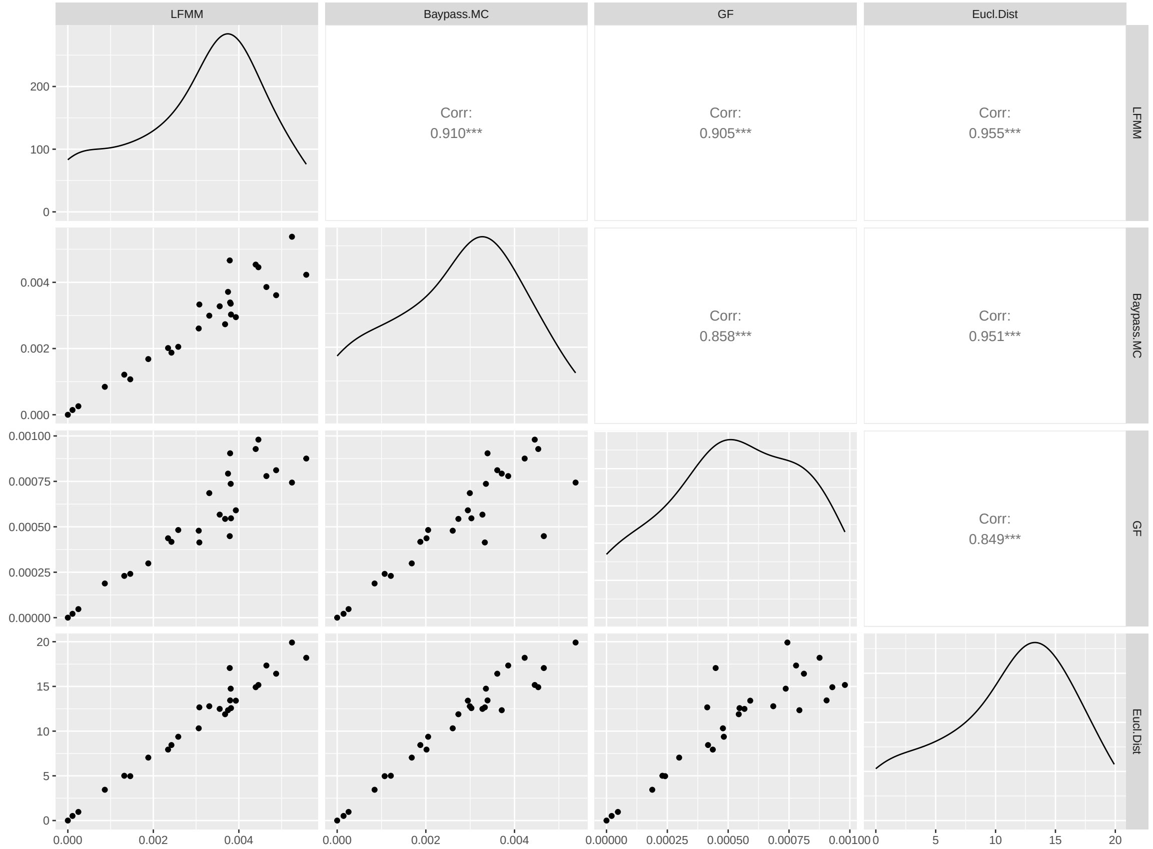


Figure S14: **Correlation between $gGO_{mc}$, $gGO_{lfmm}$, $GO_{gf}$ and Euclidean distance at the location of the 27 populations of *B. tryoni*, computed with PCs.** The population 28 is not taken into account because its GO values were not computed due to Tahiti being too far from the Australian mainland. The lower panels display the correlations between pairs of measures, while the upper panels present the Spearman correlation coefficients for each pair of GO measures. The significance of the p-value for the Spearman correlation test is denoted by stars (*** when p-value $< 0.001$), and the diagonal exhibits the distribution of each GO measure.

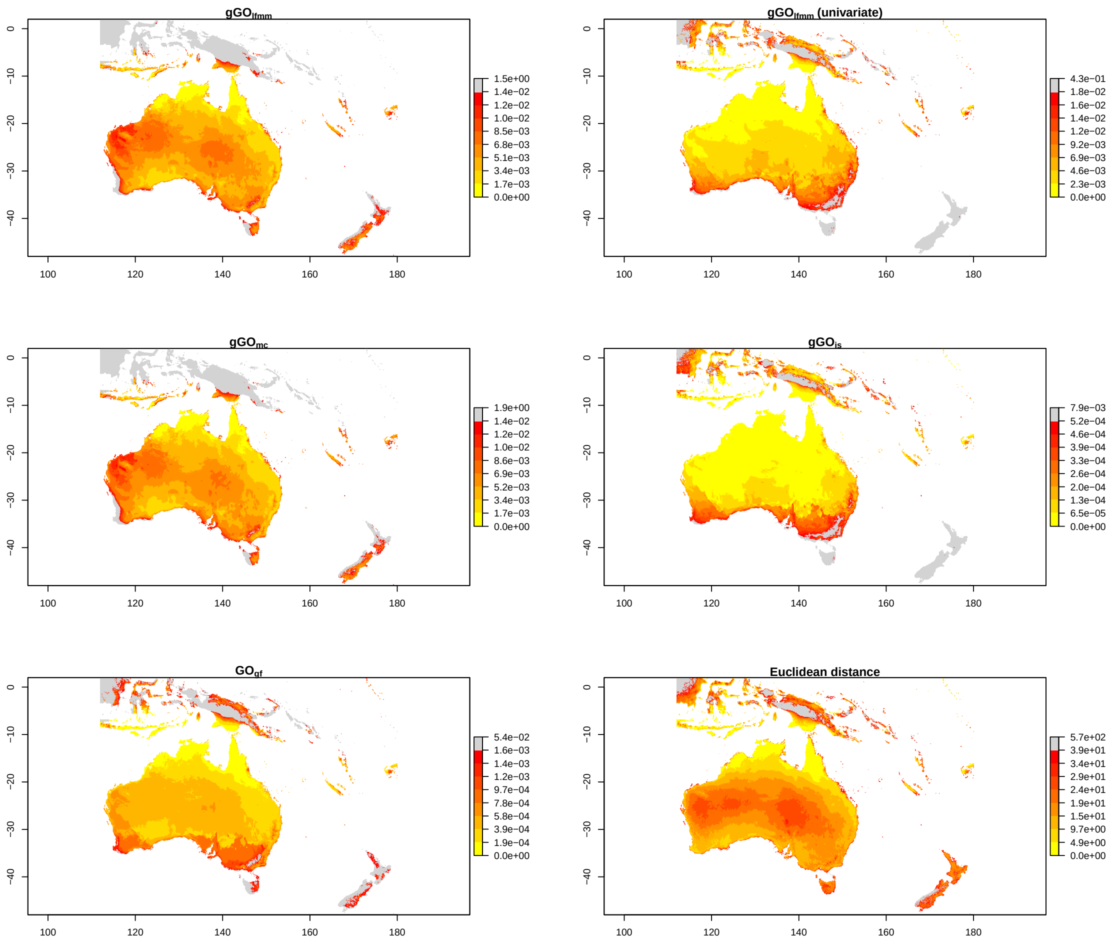


Figure S15: **GO estimated between population 1 ("source" population), and a large area in Oceania, with $gGO_{lfmm}$, $gGO_{lfmm}$ modified in ordered to treat variables independently, $gGO_{mc}$, $gGO_{is}$ and $GO_{gf}$, using six variables for GO computation (bio_3 , bio_5, bio_8, bio_9 and bio_12).** Squared Euclidean distance to the source population is also displayed. Grey pixels represent outliers values.

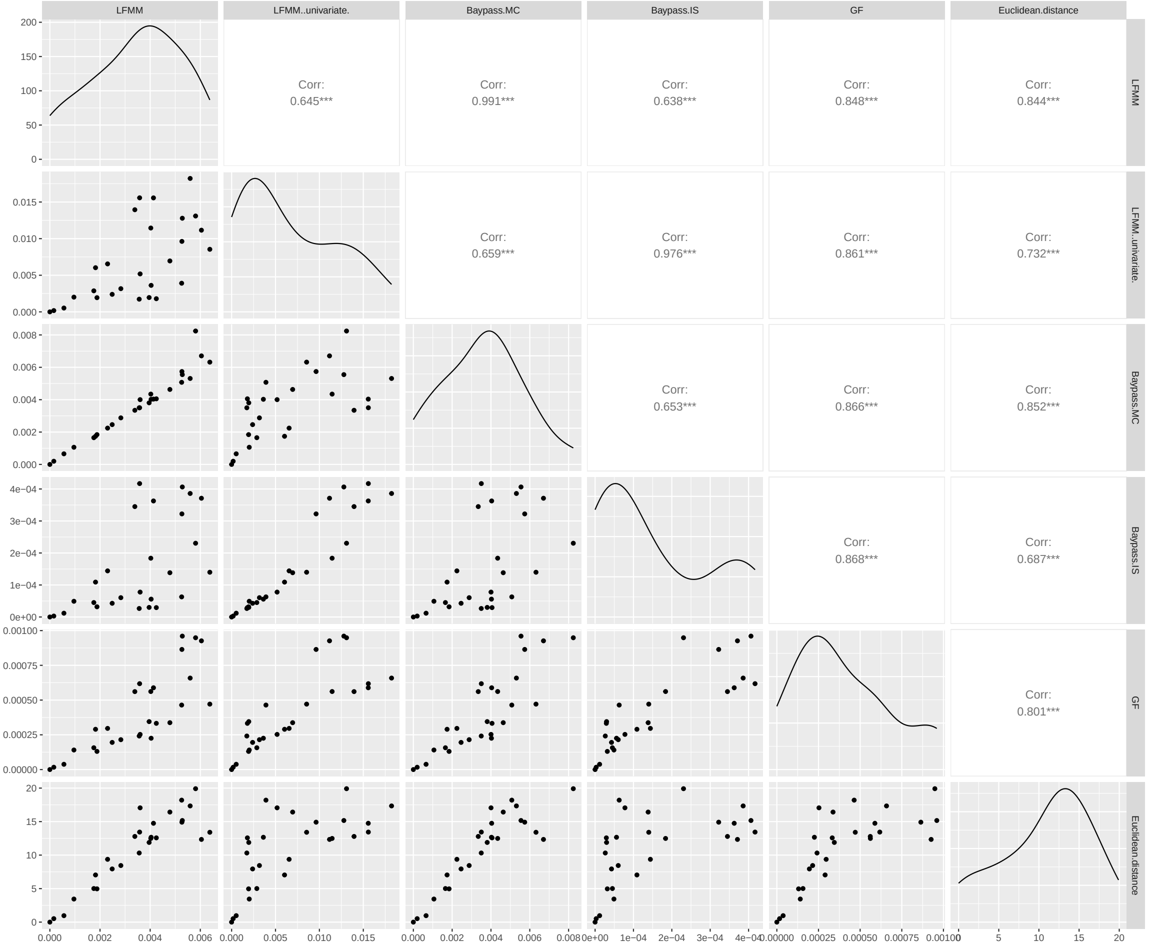


Figure S16: **Correlation between $\mathbf{gGO_{lfmm}}$, $\mathbf{gGO_{lfmm}}$ (univariate), $\mathbf{gGO_{mc}}$, $\mathbf{gGO_{is}}$, $\mathbf{GO_{gf}}$ and Euclidean distance at the location of the 27 populations of *B. tryoni*, computed on six selected variables (bio_3 , bio_5, bio_8, bio_9 and bio_12).** The population 28 is not taken into account because its GO values were not computed due to Tahiti being too far from the Australian mainland. The lower panels display the correlations between pairs of measures, while the upper panels present the Spearman correlation coefficients for each pair of GO measures. The significance of the p-value for the Spearman correlation test is denoted by stars (*** when p-value $< 0.001$), and the diagonal exhibits the distribution of each GO measure.

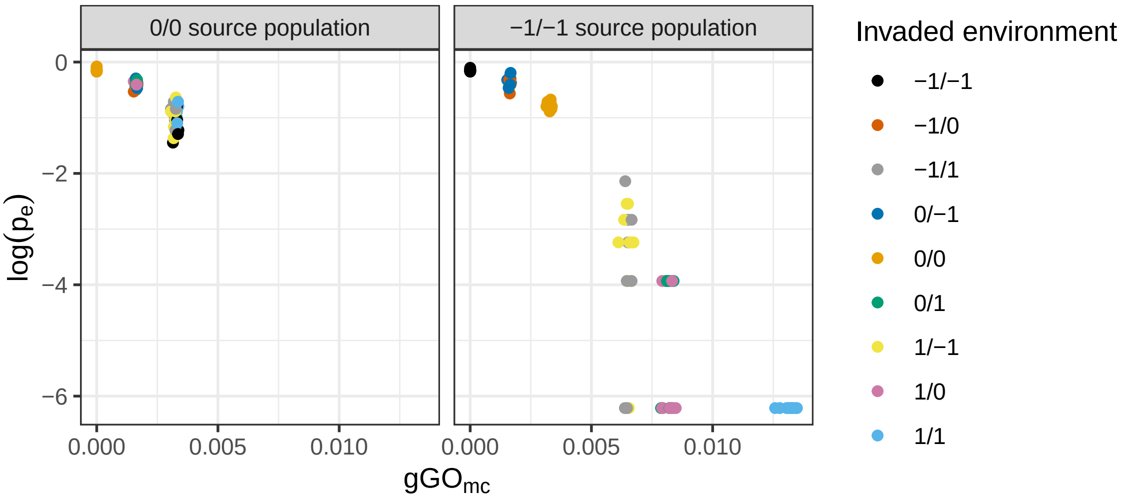


Figure S17: **Comparison between 0/0 and -1/-1 source populations concerning the correlation between GO (here $\mathbf{gGO_{mc}}$) and $\mathbf{log(}\boldsymbol{p_e}\mathbf{)}$.** On each panel, $\log(p_e)$ is plotted as a function of $\mathrm{gGO_{mc}}$ for 90 observations (10 repetitions of the native environment scenario × 9 possible invaded environments), which are combined to compute one $R^2$ value. The left panel presents results for 0/0 source population, and the right one for -1/-1 source population. In this example, migration rate is low, there are 10 invading individuals, the native environment type is L, and only causal variables were used to compute GO.

996 # B Supplementary Table

| **Migration rate** | **Environment type** | **Mean nb. of QTNs** | **Mean nb. of QTNs (MAF 1%)** | **Mean nb. of neutral mut. (MAF 1%)** | **Mean fitness** | **Mean Fst** |
|---|---|---|---|---|---|---|
| 0.005 | Linear | 1507 ± 35 | 44 ± 5 | 11 860 ± 112 | 0.952 ± 0.001 | 0.034 |
| | Mountain | 1568 ± 29 | 64 ± 10 | 11 781 ± 116 | 0.933 ± 0.002 | 0.049 |
| | Random | 1535 ± 24 | 62 ± 5 | 11 873 ± 95 | 0.945 ± 0.001 | 0.072 |
| 0.05 | Linear | 1756 ± 33 | 82 ± 12 | 11 548 ± 126 | 0.883 ± 0.002 | 0.0034 |
| | Mountain | 1955 ± 43 | 109 ± 11 | 11 504 ± 61 | 0.807 ± 0.002 | 0.0040 |
| | Random | 1852 ± 48 | 80 ± 10 | 11 509 ± 142 | 0.815 ± 0.002 | 0.0046 |

Table S1: **Information about native area simulations.** Means are computed over 10 replicates for each native area, and at the end of the 3,000 simulated generations. The standard deviation of each mean is displayed after ± sign.

### C Supplementary Text

#### C.1 Note 1: details on simulated scenarios

##### C.1.1 Fitness

The fitness of an individual $i$ living in the population $j$ will increase as the cumulative effect sizes of its QTNs (representing its phenotype $\mathrm{P}_{1,ij}$ and $\mathrm{P}_{2,ij}$) approach the values of the two environmental optima of the population $j$ at the generation $t$ ($\Theta_{1jt}$ and $\Theta_{2jt}$). Individual fitness ($\omega_{ij}$) is computed using a multivariate normal distribution with a standard deviation of $\sigma_k$ as described in Láruson *et al.* (2022) previous work :

$$\omega_{ij} = e^{\frac{-1}{2}} \left[ \left( \frac{P_{1,ij} - \Theta_{1jt}}{\sigma_k} \right)^2 + \left( \frac{P_{2,ij} - \Theta_{2jt}}{\sigma_k} \right)^2 \right] \tag{1}$$

where $\sigma_k$ reflects the intensity of stabilizing selection. Similar to the environmental optima $\Theta_{1jt}$ and $\Theta_{2jt}$, $\sigma_k$ varied along the simulation of the native area. It was initially set at 4 during the first 1000 generations, implying weak stabilizing selection, enabling high variance in individual's phenotypic values. It then gradually decreased over the next 1000 generations, ultimately reaching a value of 1.25 implying stronger stabilizing selection, which was kept constant during the final 1000 simulated generations.

##### C.1.2 Life cycle and evolutionary rates

In the native area, evolution was simulated under a Wright-Fisher model, where generations did not overlap and population size was held constant. However, reproduction was not random : the fitness of individuals corresponded to their probability of being chosen as parents for the next generation, which favored the spread of locally advantageous mutations.

In the invaded area, the simulation framework diverges from the Wright-Fisher model to allow for population extinction, essential for calculating establishment probabilities. The simulation encompasses three age classes, and includes senescence, with individuals beyond the first age class (adults) experiencing a halving of their fitness each generation. Individuals moved to the next age class when they survived, which occurred with a probability depending on their fitness. Details on life cycle are provided in Figure T1.

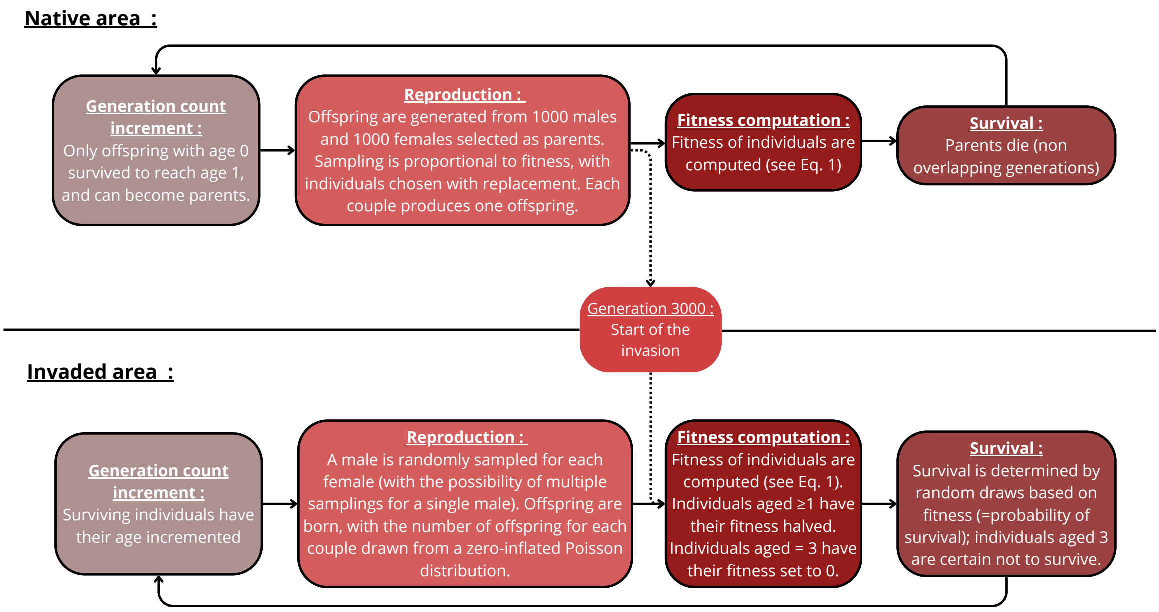


Figure T1: **Simulation life cycle in the native (top panel) and invaded area (bottom panel).**

These life cycle parameters were chosen to approximate the biology of invasive insects', with characteristics such as high fecundity, short lifespan, fast development and overlapping generation (Sakai *et al.*, 2001; Zhao *et al.*, 2023). To simulate short lifespan and fast development, the majority of individuals experience only two age classes, one as juveniles and one as adults. To account for individuals living beyond the average lifespan, their fitness is halved upon reaching the second age class. An important decrease in survival rate when population grows older has been reported in several invasive species, such as *Drosophila suzukii* (Tochen *et al.*, 2014), *B. tryoni* (Fanson and Taylor, 2012) or *Bactrocera dorsalis* (Jaleel *et al.*, 2018a).

Only adults can reproduce, with a 50% probability for a mating pair to produce offspring, considering the possibility of non-successful mating. The number of offsprings for successful mating is drawn from a Poisson distribution, with an average of 20 offspring per mating pair. This number may seem low considering that for some species such as *D. suzukii*, or *Bactrocera* species, the fecundity (=Lifetime production of offspring (eggs) by an average female that lives to the last day of life in the cohort) can sometimes reach more than 500 eggs produced (Hamby *et al.*, 2016; Jaleel *et al.*, 2018b). However, preliminary tests were conducted, exploring mean offspring numbers of 5, 20, and 100 per mating pair, and they indicated that simulation results remained relatively stable.

It's important to note that while these life cycle parameters were designed to align with the biology of insect invasive species, they can vary significantly depending on factors such as humidity, temperature, rearing conditions, population density, and more (Hamby *et al.*, 2016). Consequently, the simulations were not intended to precisely replicate the life cycle of any specific species. Instead, the primary objective was to construct a simulation model that captures the general characteristics of an insect-like species.

#### C.2 Note 2: Genetic Offset measures

##### C.2.1 GO computation

**GF correction with LFMM residuals.** LFMM are regression models which can be written as follows :

$$Y = XB^T + W + E \tag{2}$$

When studying n populations for p genetic markers, Y represents the n × p response matrix of centered allele frequencies. As explained in Caye *et al.* (2019), "the fixed effect sizes are recorded in the B matrix, which has dimension p × d", where d represents the total number of primary and nuisance variables. "The E matrix represents residual errors, and it has the same dimensions as the response matrix. The matrix W is a latent matrix of rank K, defined by K latent factors." By performing a Singular Value Decomposition (SVD) of the W matrix, one can obtain U and V matrix, corresponding respectively to the confounding factors matrix, and to the loadings of these factors :

$$W = UV^T \tag{3}$$

W matrix can then be subtracted from Y matrix as shown in equation (4), in order to obtain the residuals of the model, representing the corrected centered allele frequencies.

$$Y - W = XB^T + E \tag{4}$$

**Optimized GF.** For GF analyses, a modified version of *Gradient Forest* v 0.1.32 package was used. Indeed, this package was developed to analyze species abundance data and is not optimized for handling very large datasets such as those used in population genomics. Therefore, we have optimized the package to be able to create models using large datasets, significantly reducing computation times and memory footprint, while maintaining very similar predicted GO values (Figure T2). Note that this adapted package is (so far) compatible exclusively with continuous predictors, not with discrete ones. The resulting R function is available for download at `https://forgemia.inra.fr/simon.boitard/popgenomicprediction`

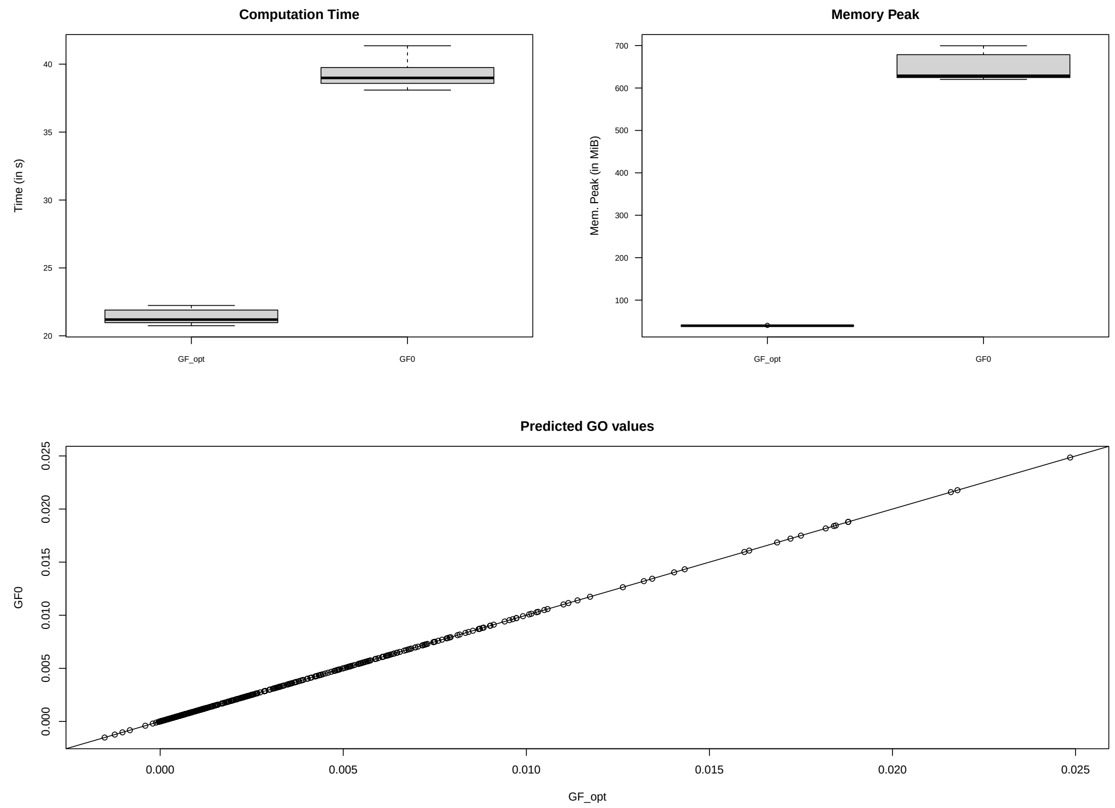


Figure T2: Comparison of performance metrics (Computation Time and Memory Use) and predicted GO values between the original Gradient Forest package (GF0) and the modified version (GF_opt). Tests were run with 10 different seeds.

#### C.3 Note 3 : Variable importances with PCs

Let $\boldsymbol{E} = \boldsymbol{UDV'}$ the singular value decomposition of the matrix of the original $D$ (centered) covariable values for the $J$ populations. By construction, the eigenvector matrix $\boldsymbol{U}$ is the scaled matrix of the $P$ principal components (PCs) (i.e., $\boldsymbol{U'U} = \boldsymbol{I}$); the diagonal matrix $\boldsymbol{D}$ contains the square root of the eigenvalues in its diagonal; and the orthogonal matrix $\boldsymbol{V}$ contains the covariable loadings of each PC (with $\boldsymbol{V'V} = \boldsymbol{I}$).

From this original decomposition, it is then easy to rescale the matrix $\boldsymbol{B_{I \times P}}$ of the regression coefficients estimated for each of the $I$ SNPs and associated to each of the $P$ PCs to obtain a $I \times D$ matrix $\boldsymbol{C}$ of regression coefficients associated to each of the original covariables. Indeed, under linear model assumptions underlying the BayPass (or LFMM) GEA models, the expectation of the vector $\boldsymbol{\alpha_i}$ of allele frequencies in the $J$ populations may be written as:

$$\mathrm{E}\left(\boldsymbol{\alpha_i}\right) = \mu_i \mathbf{1}_J + \boldsymbol{b_i'U}$$

where $\boldsymbol{b_i}$ is a $P$ length vector corresponding to the $i$th row of the regression coefficient matrix $\boldsymbol{B}$. Hence:

$$\begin{aligned}
\mathrm{E}\left(\boldsymbol{\alpha_i}\right) &= \mu_i \mathbf{1}_J + \boldsymbol{U}\boldsymbol{b_i} \\
&= \mu_i \mathbf{1}_J + \left(\boldsymbol{E}\boldsymbol{V}\boldsymbol{D^{-1}}\right)\boldsymbol{b_i} \\
&= \mu_i \mathbf{1}_J + \boldsymbol{E}\left(\boldsymbol{V}\boldsymbol{D^{-1}}\boldsymbol{b_i}\right)
\end{aligned}$$

The $D$ length vector $\boldsymbol{c_i} = \boldsymbol{V}\boldsymbol{D^{-1}}\boldsymbol{b_i}$ mays thus be interpreted as the regression coefficients associated to each original covariable. Hence the $I \times D$ matrix $\boldsymbol{C}$ can simply be obtained as:

$$\boldsymbol{C} = \left(\boldsymbol{V}\boldsymbol{D^{-1}}\boldsymbol{B'}\right)'$$

This matrix can then be decomposed to derive the importance of each covariable.
